# The *Pseudomonas aeruginosa* Membrane Histidine Kinase BqsS/CarS Directly Senses Environmental Ferrous Iron (Fe^2+^)

**DOI:** 10.1101/2025.04.06.647434

**Authors:** Alexander Paredes, Chioma Iheacho, Kelly N. Chacón, Aaron T. Smith

## Abstract

Prokaryotic two-component signal transduction systems (TCSs) are widely utilized by bacteria to respond to their environment and are typically composed of a transmembrane sensor His kinase (HK) and a cytosolic DNA-binding response regulator (RR) that work together to respond to environmental stimuli. An important TCS that regulates the expression of genes involved in biofilm formation and antibiotic resistance in many pathogens is the BqsRS/CarRS system, originally identified in *Pseudomonas aeruginosa*. Transcriptomics data suggested that the cognate *Pa*BqsRS stimulus is Fe^2+^, but *Pa*BqsS has not been characterized at the protein level, and a direct interaction between Fe^2+^ and *Pa*BqsS has not been demonstrated. In this work, we biochemically and functionally characterize intact *Pa*BqsS, an iron-sensing membrane HK, for the first time. Using bioinformatics, protein modeling, metal analyses, site-directed mutagenesis, and X-ray absorption spectroscopy (XAS), we show that *Pa*BqsS binds a single Fe^2+^ ion within its periplasmic domain containing an N/O-rich ligation sphere that includes Glu^48^ as a key metal ligand. Using activity assays, we show that both intact and truncated *Pa*BqsS have competent ATPase activities, consistent with predicted function. Importantly, we show that the ATP hydrolysis of intact *Pa*BqsS is stimulated exclusively by Fe^2+^, revealing metal-based activation of a functional, intact membrane HK for the first time. Moreover, stimulation assays of *Pa*BqsS variants demonstrate the importance of Glu^45^ and Asn^49^ in the sensing and signal transduction pathway. Taken together, this work uncovers important structural and biochemical properties that could be leveraged to target the BqsRS system for future therapeutic developments.

## INTRODUCTION

*Pseudomonas aeruginosa* is a Gram-negative, opportunistic, virulent bacterium that can survive in many conditions and is highly adaptable, making it a pressing threat to human health.^1^ As part of its adaptability, *P. aeruginosa* can grow either as a plankton (*i.e.*, freely floating in solution such as stagnate water) or within a biofilm (*i.e.*, stationary within a protective, complex, three-dimensional network composed of polysaccharides, proteins, lipids, and DNA).^1–4^ Due to its widespread nature, its adaptability, and its virulence, *P. aeruginosa* is one of the most common causes of nosocomial infections, especially for patients suffering from pneumonia, neutropenia, and/or cystic fibrosis.^1,5,6^ ^7^ Moreover, infections of *P. aeruginosa* are notoriously difficult to treat, as this bacterium continues to evolve multidrug-resistance, which is exacerbated by the ability of *P. aeruginosa* to form protective biofilms and to employ numerous virulence factors that allow this pathogen to colonize and to establish itself within a mammalian host such as a human.^8^

An important factor that governs the infectivity of many pathogens, such as *P. aeruginosa*, is the ability of these bacteria to acquire key nutrients from their hosts, including essential transition metal ions. Elements such as Mn, Fe, Cu, and Zn are generally necessary for the normal metabolic functions of virtually all bacteria, and their limited presence can often control microbial virulence (Fig. 1).^9–12^ However, these crucial metal ions can also be toxic to cells at high concentrations, as they can disrupt cellular functions through protein and/or nucleic acid misfolding and via the generation of reactive oxygen and nitrogen species (ROS and RNS). Due to their dual nature, bacteria have developed complex mechanisms to sense the presence of transition metals and to regulate their influx and efflux, especially at the host-pathogen interface.^10,12–16^

**Figure 1.**
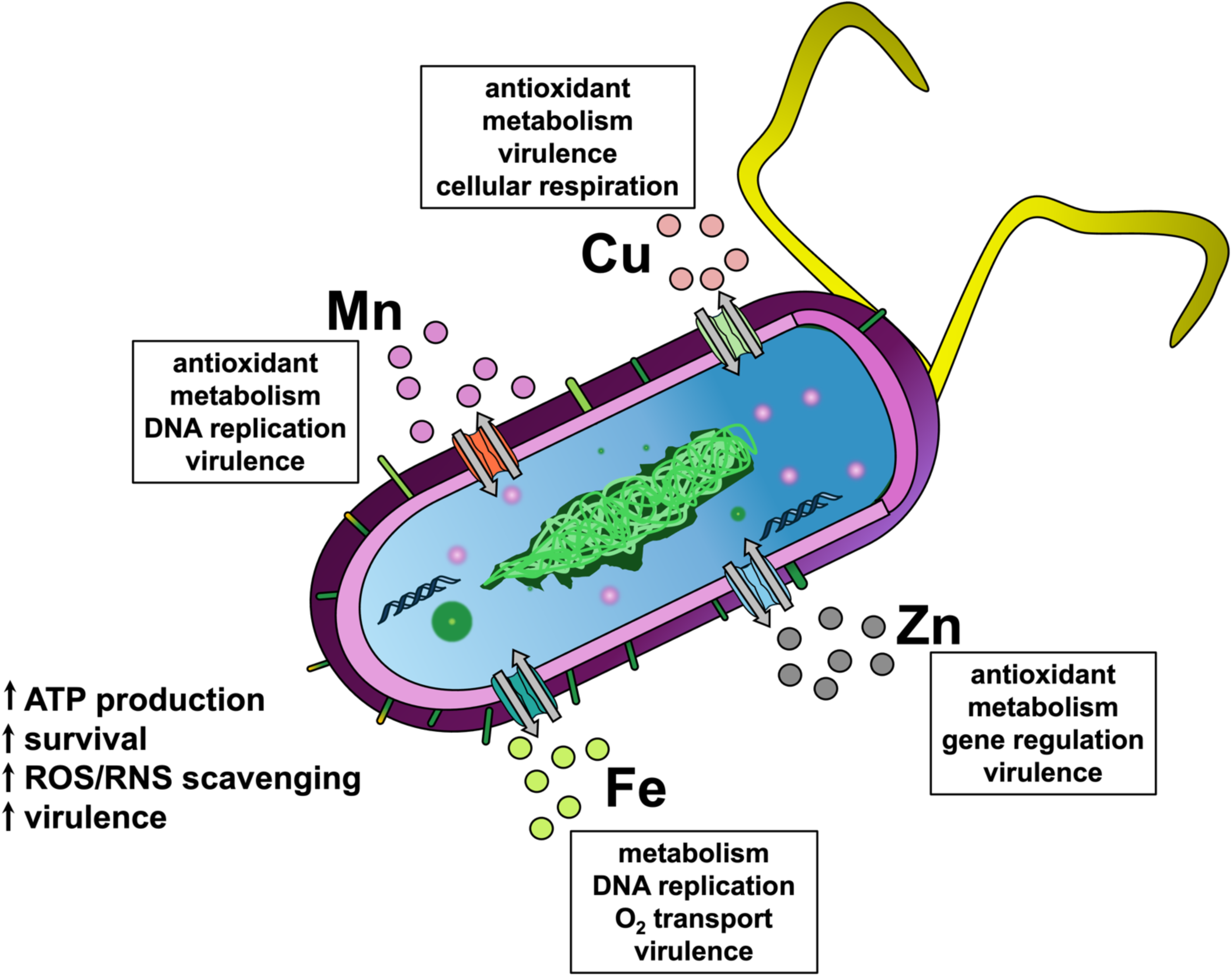
Transition metals such as Mn, Fe, Cu, and Zn are critical for normal bacterial cellular function and may be leveraged by pathogenic strains. Transition metals are involved in several critical metabolic roles within prokaryotes, such as electron transport, DNA replication, and even transcriptional regulation. Importantly, transition metals are also a part of microbial virulence, as these micronutrients can help increase ATP production, can be leveraged to defend against ROS/RNS, and can even be involved in biofilm formation and dispersion.

Two-component signal transduction systems (TCSs) are common mechanisms used by bacteria to sense the presence of environmental nutrients, such as transition metals, and these systems function to regulate intracellular concentrations of metals, allowing bacteria to adapt to a wide array of conditions found within the human host.^5,9,17–19^ A bacterial TCS is canonically composed of two separate proteins: a transmembrane sensor His kinase (HK) and a cytosolic DNA-binding response regulator (RR).^20,21^ The extracytoplasmic domain of the HK senses the external nutrient/stimulus, leading to an autophosphorylation event that occurs at the HK cytoplasmic domain. This domain consists of two sub-domains spatially adjacent to one another: the catalytic ATP-binding (CA) domain that catalyzes the transfer of the γ-phosphate from ATP to the HK, and the dimerization and His phosphotransfer (DHp) domain that houses a conserved phosphate-accepting His residue. Once phosphorylated, this labile moiety is then transferred from the cytoplasmic domain of the HK to a conserved Asp residue on the cytoplasmic RR. Phosphorylation of the RR leads to dynamical changes, often accompanied by protein dimerization, that increase the affinity of the RR towards its specific DNA sequence(s), ultimately resulting in the transcriptional regulation of target genes.^20,21^ The types of stimuli that may be sensed are often specific (*e.g.*, H^+^, transition metals, small molecules), while the genes targeted by the RR can be varied and often remodel the metabolic architecture of the cell, frequently increasing the infectivity of pathogenic bacteria through the expression of numerous virulence factors.^20–28^

BqsRS (also known as CarRS) is an important but understudied metal-sensing TCS first discovered in *P. aeruginosa* that regulates biofilm formation and decay.^29,30^ Like most canonical TCSs, BqsRS is composed of a membrane-bound HK, BqsS, and its cognate RR, BqsR.^29,30^ Initial transcriptional profiling showed that the membrane HK BqsS specifically recognizes Fe^2+^ at μM concentrations,^30^ while subsequent studies have shown that BqsRS may also respond to mM concentrations of Ca^2+^.^31^ These metal concentrations are similar to levels present in the cystic fibrosis lung environment,^32–37^ and *bqsS* transcripts have been detected in cystic fibrosis sputum samples.^38^ Further underscoring its importance, functional studies in *P. aeruginosa* have linked the presence of BqsRS to the regulation of genes involved in biofilm formation and antibiotic resistance, such as those responsible for rhamnolipid production and the formation of short chain quorum sensing signals like *N*-butyryl-*L*-homoserine lactone.^29^ *Pa*BqsS has been proposed to sense Fe^2+^ via a conserved ExxE motif present in the BqsS extracytoplasmic domain, but no metal-protein interaction in this system has been directly observed. In fact, to our knowledge no previous work has shown a stable interaction between any transition metal and any intact membrane HK, and how this important system selects for Fe^2+^ vs Ca^2+^ is unclear, prohibiting targeting of this TCS to treat *P. aeruginosa* infections.

In this work, we biochemically and biophysically characterize the metal-binding and functional properties of the *P. aeruginosa* membrane HK BqsS in its intact and truncated forms. Recombinantly expressed, detergent-solubilized *Pa*BqsS is purified to homogeneity and is consistent with a dimeric oligomerization, similar to other canonical HKs. Importantly, we quantify and characterize a stable interaction between Fe^2+^ and intact *Pa*BqsS for the first time. Using X-ray absorption spectroscopy (XAS), we show that Fe^2+^ binds to the intact membrane HK in an octahedral geometry, coordinated by 6 N/O metal ligands. Using site-directed mutagenesis and XAS, we show that Glu^48^ is critical for Fe^2+^ binding, while Glu^45^ is implicated in maintaining the correct Fe^2+^ coordination geometry. Employing enzymatic assays, we characterize the ATP hydrolysis of intact and variant *Pa*BqsS in the presence and absence of Fe^2+^ to corroborate the critical residues involved in Fe^2+^ coordination and to define important auxiliary amino acids involved in signal transduction. To our knowledge, this study represents the first characterization of an intact HK bound to, and stimulated by, its cognate metal. Moreover, results from this work define key structural and biochemical properties that are important for the sensing, coordination, and signal transduction of Fe^2+^ by the *Pa*BqsRS two-component system.

## RESULTS

### Expression and purification of intact, homogenous PaBqsS

To explore the biophysical properties of intact *Pa*BqsS, we first cloned, expressed in *Escherichia coli*, isolated, solubilized, and purified intact *Pa*BqsS to homogeneity. Surprisingly, very few intact HKs have been purified, and no intact metal-sensing HK has been biophysically characterized in the presence of its cognate metal stimulus, likely due to the difficulty of producing and purifying the full-length membrane protein.^39–42^ After cloning intact *Pa*BqsS comprising its periplasmic sensor domain, two transmembrane helices, and a soluble domain CA domain linked to a DHp domain (Figs. 2a,2b), we verified its overexpression and optimized its membrane isolation conditions. We then tested multiple different non-denaturing detergents for their ability to solubilize *Pa*BqsS stably (Fig. S1). Based on western blotting and downstream characterization, we settled on the use of *n*-tetradecylphosphocholine (Fos-Chol-14) as the detergent for large-scale purification, tag cleavage, and gel filtration studies, as other detergents such as *n*-dodecyl-β-D-maltoside (DDM) and lauryldimethylamine-*N*-oxide (LDAO) caused irreversible protein aggregation, while our *Pa*BqsS isolated in Fos-Chol-14 was highly pure and homogeneous (Fig. 2b). Size-exclusion chromatography indicates that *Pa*BqsS migrates as a dimer in two different detergents (Fos-Chol-14 and glyco-diosgenin (GDN); Fig. 2b and Fig. S2), which was further confirmed by rapid dilution coupled with mass photometry in Fos-Chol-14 (Fig. S3). These observations are consistent with previous studies that have shown that most canonical membrane HKs exists as homodimers^43,44^ and with a high-confidence AlphaFold model that we were able to generate (Fig. 2b, Fig. S4). Moreover, the interaction of *Pa*BqsS with its cognate stimulating ligand (Fe^2+^) also suggests an operative dimer is in solution (*vide infra*). Thus, we were able to isolate and to purify intact, dimeric *Pa*BqsS for the first time for downstream biophysical and enzymatic characterizations.

**Figure 2.**
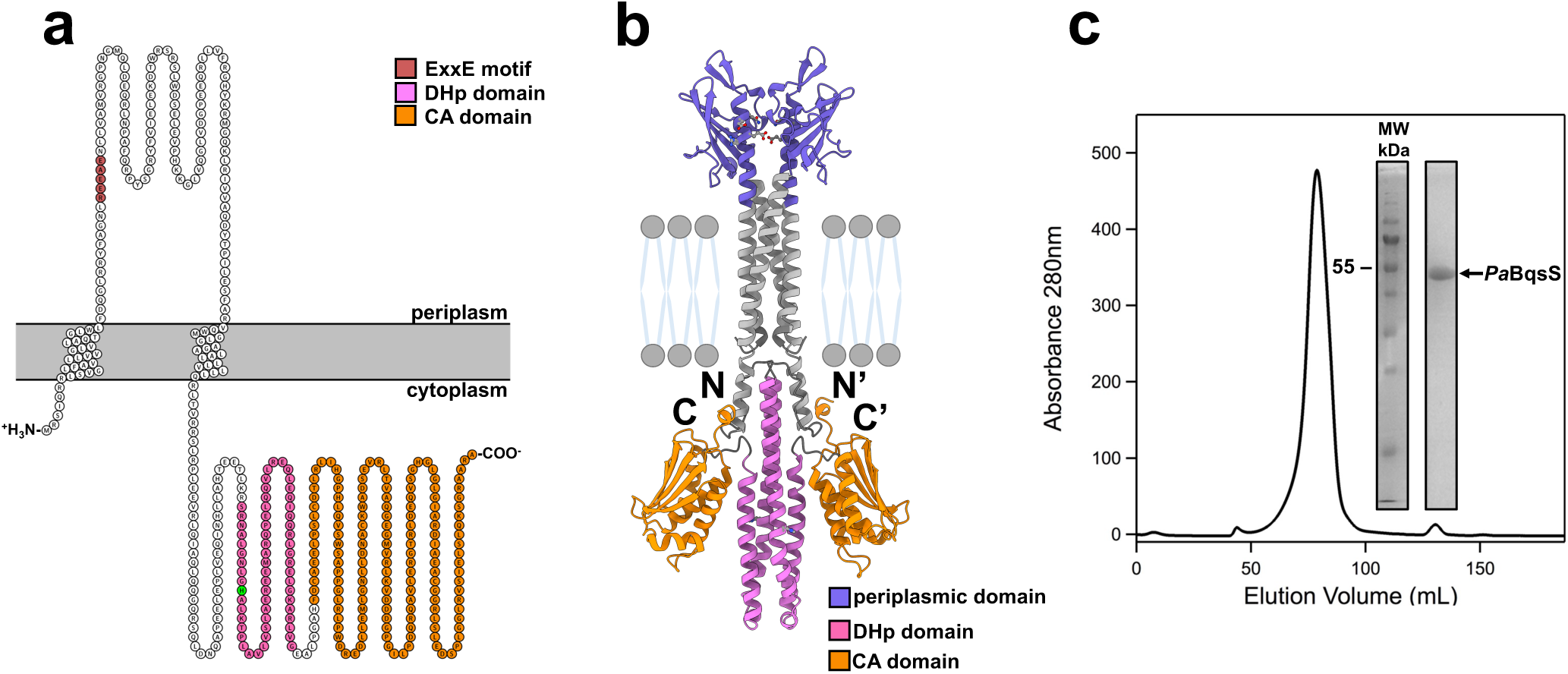
Topological modeling and purification to homogeneity of intact *Pa*BqsS. **a**. A cartoon image of the predicted topology of *Pa*BqsS based on the TMHMM server.^81,82^ The predicted topology of intact *Pa*BqsS includes two transmembrane helices, a periplasmic domain with a putative iron-sensing ExxE motif, and a long C-terminal domain composed of CA and DHp sub-domains. Image partially created using Protter.^83^ **b**. The AlphaFold model of intact, dimeric *Pa*BqsS with the various domains (periplasmic, DHp, CA) color-coded. **c**. The size-exclusion chromatogram (SEC) and SDS-PAGE analysis (*inset*) of cleaved, purified *Pa*BqsS in Fos-Choline-14 demonstrates good homogeneity and purity when recombinantly produced in *E. coli*.

### PaBqsS binds Fe^2+^ at a dimeric interface comprising an octahedral, N/O-rich ligation sphere

A previous *in vivo* study proposed that *Pa*BqsS binds Fe^2+^ by utilizing an ExxE motif in its HK periplasmic domain,^45^ but the atomic level details and stoichiometry of this process were not defined. We first wondered whether studies on other iron homeostasis proteins would shed light onto how *Pa*BqsS binds iron. An ExxE motif has been implicated in iron transport both in bacteria (*e.g.*, FeoB)^46^ and yeast (*e.g.*, *Saccharomyces cerevisiae* and *Candida albicans* FTR proteins)^47,48^, as well as in iron storage (*e.g.*, ferritin)^49^, although the selectivity of this motif for Fe^3+^ *versus* Fe^2+^ is unclear. We then considered if this motif were conserved amongst putative iron-sensing HKs in general. We compiled the evolutionarily divergent sequences of the few iron-sensing HKs that have been characterized at either the *in vitro* or the *in vivo* levels in the literature (*Pa*BqsS, *Ec*BasS, *Xanthomonas campestris* VgrS, *Klebsiella pneumoniae* PmrB, and *Haemophilus influenzae* FirS),^30,50–54^ we used TMHMM to predict the extent of the periplasmic domains in each, and we searched these domain sequences for the presence of an ExxE motif (Fig. 3a). In almost all cases, an ExxE motif was predicted to be present in the HK periplasmic domain, often preceded by a basic residue (Arg or His), and often proceeded by an amide residue (Asn or Gln) or an indole residue (Trp), with the exception of *Hi*FirS that has a slightly longer ExxxE motif (Fig. 3a). We then used AlphaFold modeling to predict the dimeric structures of these domains, with the hypothesis that these structural predictions would provide insight into metal-ligand coordination and stoichiometry. Interestingly, several of the AlphaFold periplasmic domain models such as *Pa*BqsS, *Ec*BasS, and *Kp*PmrB appear to place these ExxE motifs at the dimer interface (common to the binding interface for many HKs),^43,44^ whereas the AlphaFold periplasmic domain models of *Xc*VgrS and *Hi*FirS place the Exx(x)E motifs distal to the dimeric interface (Fig. 3b-f). Whether these motifs exclusively bind Fe^2+^, Fe^3+^, or both is unclear, but the flanking residues could be involved in this selectivity. Moreover, whether the ExxE motif is clustered at, or distant from, the dimeric interface may implicate different binding stoichiometries.

**Figure 3.**
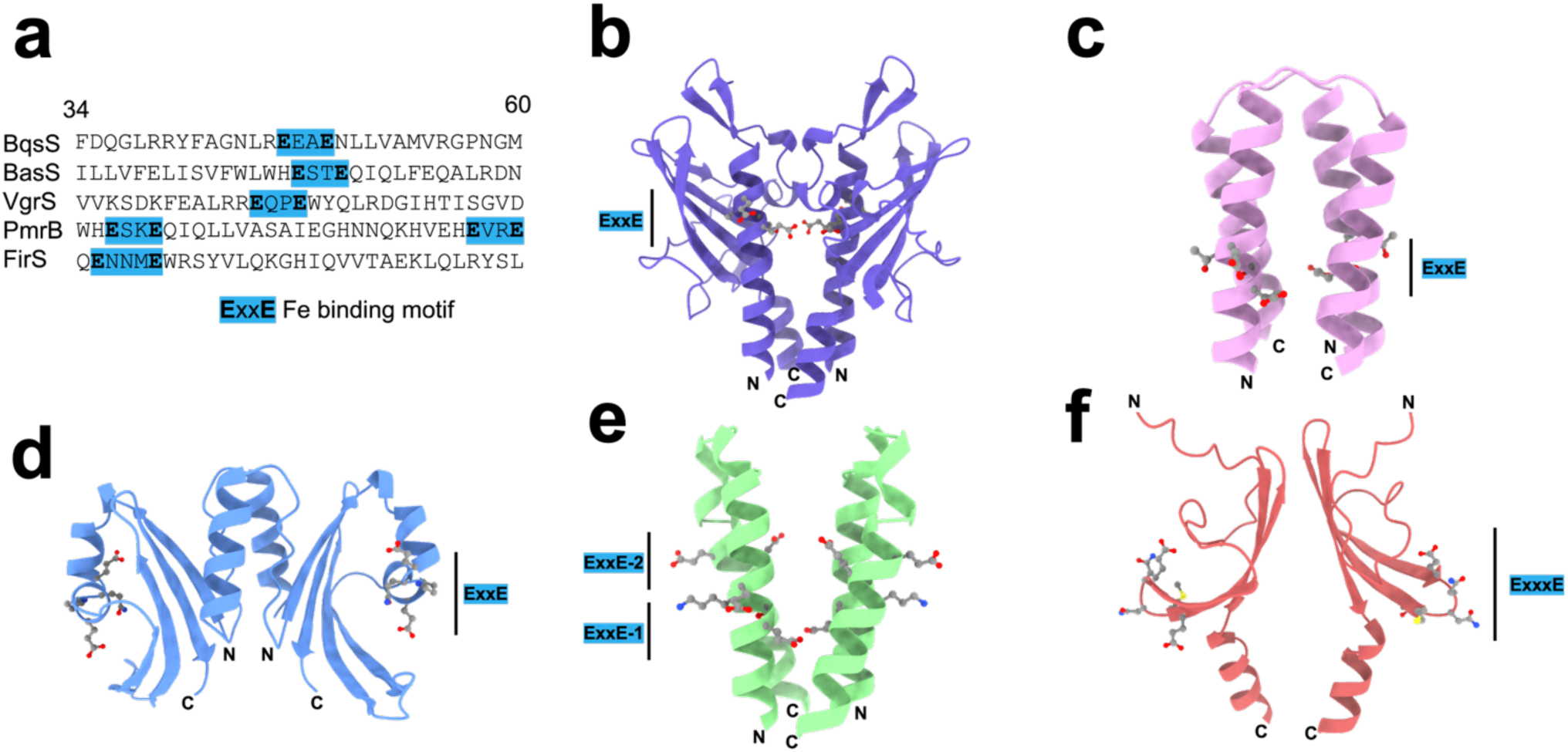
A periplasmic Exx(x)E motif is conserved across dimeric, iron-sensing HKs. **a**. Partial periplasmic amino acid sequences illustrate the presence of the Exx(x)E motif in several iron-sensing HKs. While the general Exx(x)E motif is present in all iron-sensing HKs, its internal and flanking residues differ, as does its predicted structural positioning relative to the dimer interface. The numbers at the top of the sequences indicate the positioning relative to *Pa*BqsS. **b**. The AlphaFold prediction of the periplasmic domain *Pa*BqsS suggests an ExxE motif is present at the dimeric interface. **c**. The AlphaFold prediction of the periplasmic domain of *E. coli* BasS, an iron-sensing HK, suggests an ExxE motif is present at the dimeric interface. **d**. The AlphaFold prediction of the periplasmic domain of *Xanthomonas campestris* VgrS, an iron-sensing HK, suggests an ExxE motif is present but is distal to the dimeric interface. **e**. The AlphaFold prediction of the periplasmic domain of *Klebsiella pneumoniae* PmrB, and iron-sensing HK, suggests two ExxE motifs are present at the dimeric interface. **f**. The AlphaFold prediction of the periplasmic domain of *Haemophilus influenzae* FirS, an iron-sensing HK, suggests a longer ExxxE motif is present but is distal to the dimeric interface.

We then tested whether *Pa*BqsS would bind to Fe^2+^ in a stable manner, and we biophysically characterized this binding interaction and compared it to our AlphaFold model (Fig. 2b, Fig. S4). We focused specifically on Fe^2+^ and not Fe^3+^ as previous research has shown that *Pa*BqsS has no transcriptional response to Fe^3+^ compared to Fe^2+^;^45^ moreover, free Fe^3+^ is highly insoluble and is thus bound tightly to small molecules and proteins *in vivo*, making uncomplexed Fe^3+^ an unlikely signal for an iron-responsive HK. We then titrated purified, intact *Pa*BqsS with excess Fe^2+^ under anoxic conditions, buffer exchanged copiously, and measured *ca.* 0.5 mol. eq. of Fe^2+^ (0.50 ± 0.06) bound per polypeptide using the ferrozine assay, a spectrophotometric assay to detect protein-bound Fe^2+^ (Fig. 4a).^55,56^ As sensor HKs largely function as homodimers^43,44^, as our gel filtration data and AlphaFold model suggest that *Pa*BqsS is also a dimer (Fig. 2b,c), and as our modeled metal-binding site appears to be at the homodimer interface (Fig. 3b), we initially hypothesized that a single Fe^2+^ ion binds per *Pa*BqsS dimer with each E^45^xxE^48^ motif contributing to binding. However, we could not initially rule out that Asn^49^ (directly following Glu^48^) might also contribute to Fe^2+^ binding. To test this hypothesis, we used site-directed mutagenesis and created several *Pa*BqsS variants to express and to purify: E45A, E48A, E45A/E48A, and N49A. While we could purify each of the single, intact *Pa*BqsS variants (Fig. S5), the double variant (E45A/E48A) failed to express and to purify, suggesting that the protein may misfold when both Glu residues are lost. Interestingly, only one modification (E48A) appeared to affect Fe^2+^ binding consequentially, as both E45A and N49A bound a similar stoichiometry of Fe^2+^ per polypeptide compared to WT *Pa*BqsS based on ferrozine assays, while E48A *Pa*BqsS failed to bind Fe^2+^ appreciably (Fig. 4a). These data indicate that Glu^48^ is a critical ligand in the stable coordination of Fe^2+^, while Glu^45^ and/or Asn^49^ are less critical, at least in initial Fe^2+^ coordination.

**Figure 4.**
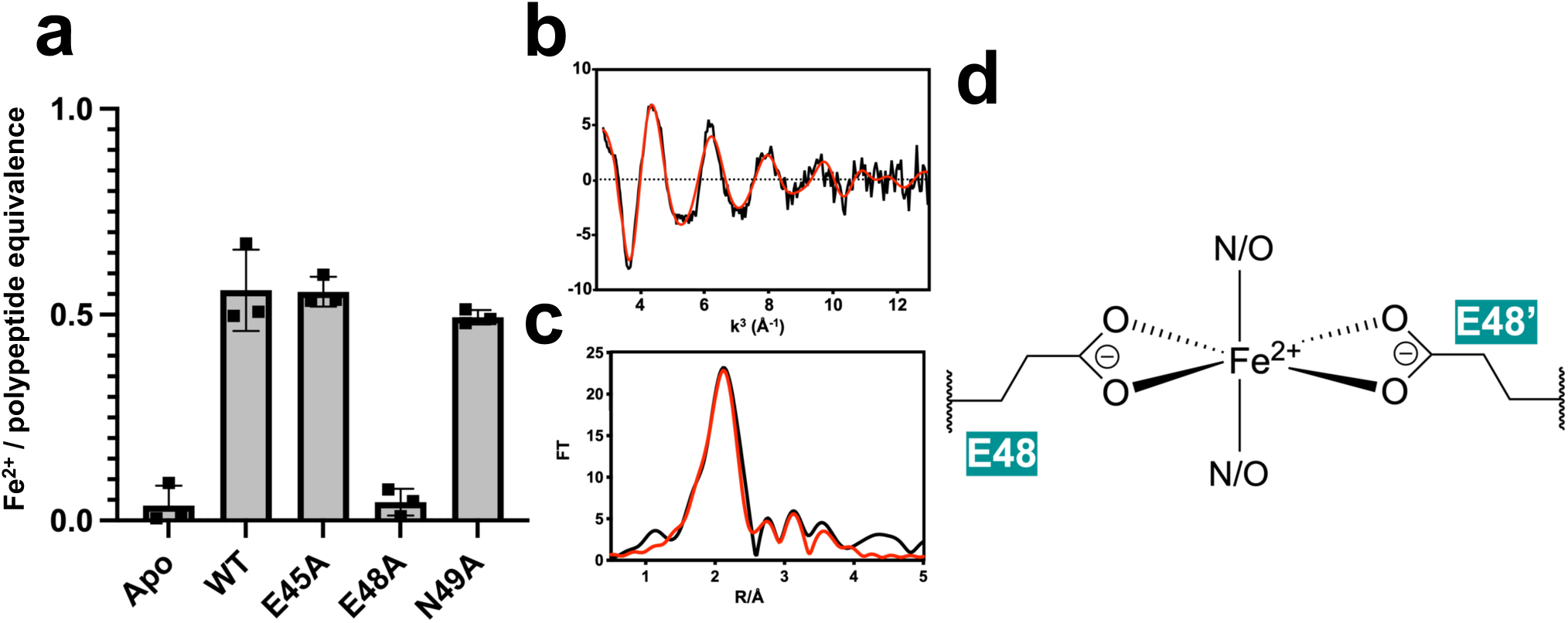
Intact *Pa*BqsS binds Fe^2+^ at its dimeric interface using an octahedral N/O motif with Glu^48^ a key metal ligand. **a**. WT, intact *Pa*BqsS binds approximately one Fe^2+^ ion per dimer, and the loss of Glu^48^ but not the loss of Glu^45^ nor the loss of Asn^49^ disrupted Fe^2+^ binding. All values were determined in triplicate (represented by each point), and the error bar represents one standard deviation of the mean. **b,c**. EXAFS of *Pa*BqsS bound to Fe^2+^ under anoxic conditions. The processed EXAFS (**b**) and resulting Fourier transformed data (**c**) are consistent with the presence of Fe^2+^ in a six-ligand geometry composed of an N/O-rich environment. The black traces represent the experimental data, while the red traces represent the EXCURVE-fitted data. **d**. Cartoon representation of the proposed Fe^2+^ coordination within the periplasmic domain of *Pa*BqsS with Glu^48^ representing a key Fe^2+^ ligand.

To corroborate these mutagenesis data and to gain greater insight into the identity of the Fe^2+^ ligands and the structure of the Fe^2+^ binding site, we used X-ray absorption spectroscopy (XAS) (Fig. 4b,c), a powerful technique that is capable of probing the oxidation state, coordination number, coordination geometry, and ligand identies of metals bound to biological macromolecules.^55^ To do so, we first anoxically prepared large concentrations (>1 mM) of Fe^2+^-bound WT *Pa*BqsS and analyzed it using XAS (Fig. 4b,c). The energy of the inflection of the X-ray absorption near-edge structure (XANES) spectrum of Fe^2+^-bound WT *Pa*BqsS as well as its weak pre-edge features are consistent with an octahedrally-coordinated Fe^2+^ ion (Fig. S6). Simulations of the extended X-ray absorption fine structure (EXAFS) data of *Pa*BqsS (Fig. 4b,c) fitted based on small-molecule complexes^56^ reveal an N/O-based primary shell environment surrounding the Fe^2+^ ion, with 4 shorter N/O scatterers at 2.04 Å from the Fe^2+^ ion, 2 longer N/O scatterers at 2.14 Å from the Fe^2+^ ion, and strong carbon secondary shell contributions from the C-bonded N/O ligands (Table 1). Based on our mutagenesis results (Fig. 4a), we believe that the 4 shorter N/O ligands to Fe^2+^ at 2.04 Å are due to tight interactions between Fe^2+^ and Glu^48^. However, we could not initially rule out whether Glu^45^ or Asn^49^ occupied the 2 longer N/O ligands to Fe^2+^ at 2.14 Å, so we anoxically prepared large concentrations (>1 mM) of the Fe^2+^-bound E45A and N49A variants for XAS. Unfortunately, at high concentrations we encountered stability issues with the E45A variant that preclude high-resolution and high-confidence EXAFS modeling; however, the XANES spectrum of Fe^2+^-bound E45A *Pa*BqsS displays a distinct, increased pre-edge feature strongly suggesting a change in coordination geometry for this protein variant (Fig. S6). In contrast, the XANES spectrum of Fe^2+^-bound N49A *Pa*BqsS is nearly identical to the WT protein (Fig. S6) and its experimental and fitted EXAFS data suggest a nearly identical ligation sphere in the N49A variant compared to the WT protein (Fig. S7, Table S1). Thus, we believe that the 2 longer and weaker Fe^2+^-N/O ligands at 2.14 Å are likely from a neutral Glu^45^ sidechain, but we cannot rule out other neutral N/O ligands. To our knowledge, this discovery represents the first time that the coordination site of Fe^2+^ bound to an intact, iron-sensing HK has been characterized.

**Table 1.** Fits obtained for the Fe K-EXAFS of WT *Pa*BqsS bound to Fe^2+^. Fits were determined by curve fitting using the program EXCURVE (version 9.2).

| Sample | Fit index<sup>a</sup> | Fe-O/N (non His) |  |  | Fe-C (non His) |  |  | E<sub>o</sub><sup>e</sup> (eV) |
| --- | --- | --- | --- | --- | --- | --- | --- | --- |
|  |  | No<sup>b</sup> | R<sup>c</sup> (Å) | DW<sup>d</sup> (Å<sup>2</sup>) | No | R (Å) | DW (Å<sup>2</sup>) |  |
| WT *PaBqsS* + Fe<sup>2+</sup> | 0.076 | 2.0 | 2.14 | 0.008 | 2.0 | 2.89 | 0.013 | -4.77 |
|  |  | 4.0 | 2.04 | 0.022 | 2.0 | 3.09 | 0.004 |  |
|  |  |  |  |  | 2.0 | 3.70 | 0.004 |  |

<sup>a</sup>The least-squares fitting parameter (see *Methods*) <sup>b</sup>Coordination number <sup>c</sup>Bond length <sup>d</sup>Debye-Waller factor <sup>e</sup>Photoelectron energy threshold

### Binding of Fe^2+^ to the PaBqsS HK specifically and significantly stimulates ATPase activity

The AlphaFold model of intact *Pa*BqsS (Fig. 2b, Fig. S4) suggests that binding Fe^2+^ in the periplasmic sensor domain of this HK may have downstream functional consequences. Specifically, as we have shown that a single Fe^2+^ ion binds within the periplasmic sensor domain utilizing Glu^48^ and, likely Glu^45^, we considered how this binding event could alter *Pa*BqsS function. The entire stretch of the E^45^xxE^48^ sequence is predicted confidently to be located along an α-helix, with Glu^45^ and Glu^48^ from each protomer suggested to be proximal (*ca*. 6 Å) to one another (Fig. 5; Fig. S4). However, in our AlphaFold model each of these sets of ligands on each protomer are modeled distal to the other set with *ca*. 15 Å being the shortest distance from Glu^45^ on protomer 1 to Glu^45^ and protomer 2 (Fig. 5). If accurate, this distance would necessitate a large conformational change in order to facilitate binding of Fe^2+^ at the dimeric interface of the HK. Such a conformational change may seem extreme, but research has shown that a structural reorientation is important for sensor domains of TCSs,^57^ and such movements has even been structurally observed for other stimulus-sensing membrane HKs.^42^

**Figure 5.**
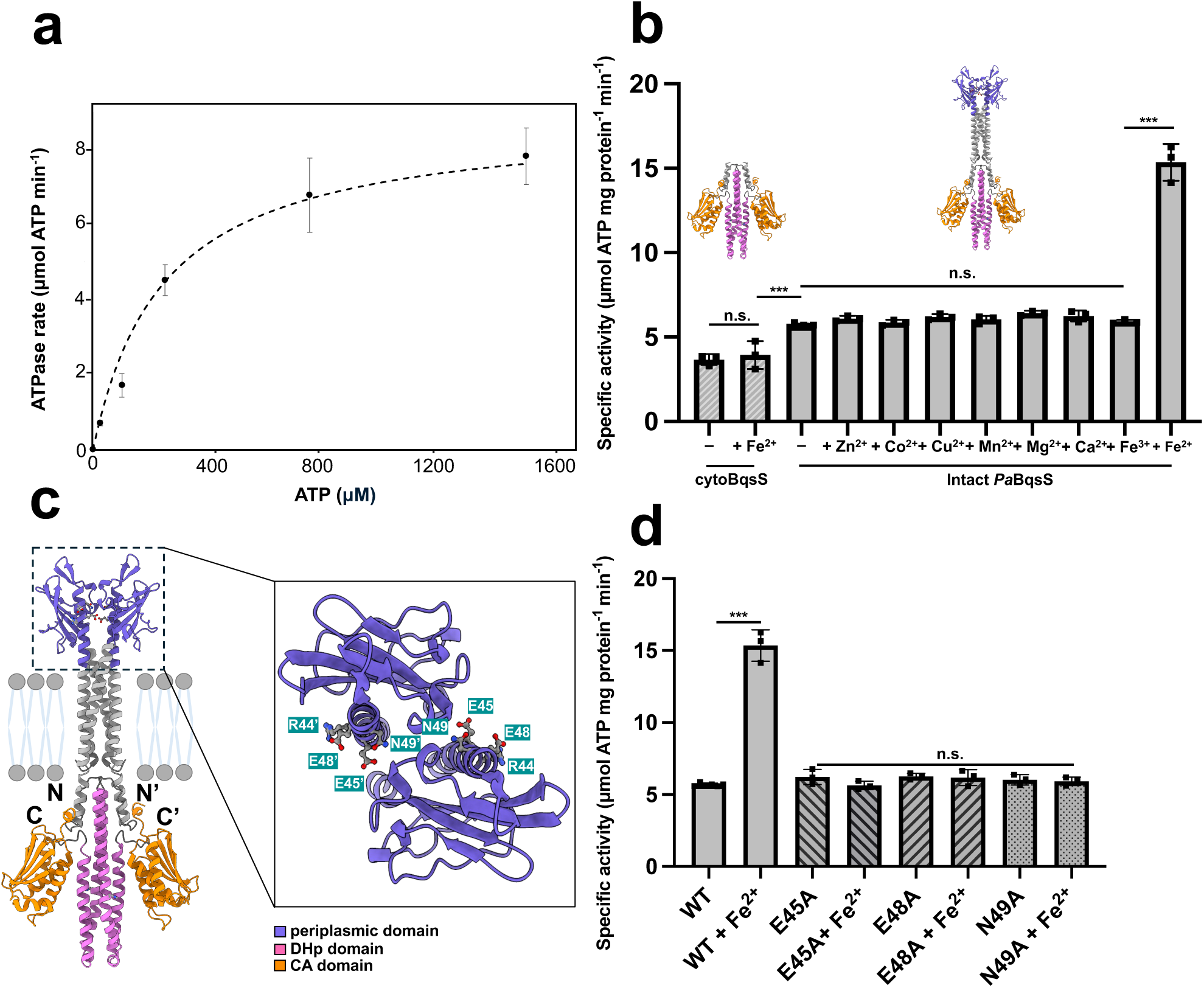
The binding of Fe^2+^ in the periplasmic sensor domain of *Pa*BqsS is transmitted through the transmembrane region to the DHp and CA domains of *Pa*BqsS leading to an alteration in ATP hydrolysis rates. **a**. The Michaelis-Menten profile of cytoBqsS-catalyzed ATP hydrolysis demonstrates a maximum velocity (*V_max_*_)_ of 8.82 ± 0.42 μmol ATP*×*min^-1^ and a Michaelis constant (K_M_) of 244.34 ± 17.28 μM. **b**. The specific ATPase activity of cytoBqsS and intact *Pa*BqsS both with and without the presence of divalent metals. There is no difference in ATPase activity in the presence of metals for cytoBqsS, while intact *Pa*BqsS has an Fe^2+^-dependent increase in ATPase activity. n.s. indicates not statistically significant (*P* > 0.05) whereas *** indicates *P* < 0.001 based on the student’s t-test. **c**. The AlphaFold model of intact *Pa*BqsS with the various domains (periplasmic, DHp, CA) color-coded. *Inset*. A conformational change in the periplasmic sensor domain may be needed to coordinate Fe^2+^, which could be transmitted through the transmembrane region to the intracytoplasmic domains. The amino acids comprising the R^44^E^45^xxE^48^N^49^ sequence in the periplasmic sensor domain are highlighted. **d**. The specific ATPase activities of intact and variant *Pa*BqsS indicate that E45, E48, and N49 play key roles in metal binding, signal transduction, and activity. All values were determined in triplicate (represented by each point), and the error bar represents one standard deviation of the mean.

Alterations along these α-helices that span across the membrane (*i.e.*, from extracytoplasmic to intracytoplasmic) are hypothesized to propagate to the intracellular DHp and CA domains to alter HK function and ATP hydrolysis, which we then tested. First, to separate any potential effects of the presence of Fe^2+^ on ATP hydrolysis catalyzed by just the intracellular, cytosolic DHp and CA domains, we cloned, expressed, and purified a construct corresponding to only this region (cytoBqsS; amino acids 210-438) (Fig. S8). Because sensor HKs all contain a CA domain that catalyzes ATP-dependent protein kinase activity,^21^ we used a modified malachite green assay^58^ to measure the ATPase activity of *Pa*BqsS. Indeed, isolated cytoBqsS is a competent ATPase with a maximal velocity (*V_max_*_)_ of 8.82 ± 0.42 μmol ATP×min^-1^ (Fig. 5a), a Michaelis constant (K_M)_ of 244.34 ± 17.28 μM (Fig. 5a), and a specific activity of 3.64 ± 0.64 μmol ATP×mg protein^-1^×min^-1^ (Fig. 5b). While cytoBqsS also bears an ExxE motif, it failed to bind Fe^2+^ and, consistent with this notion, addition of Fe^2+^ to cytoBqsS showed no effect on specific activity (Fig. 5b; 3.93 ± 0.82 μmol ATP×mg protein^-1^×min^-1^, *P* > 0.05). We then tested the ATPase activity of intact *Pa*BqsS purified in Fos-Chol-14 and showed that it was indeed active as well, with specific activity of 5.64 ± 0.38 μmol ATP×mg protein^-1^×min^-1^, modestly higher than cytoBqsS alone (Fig. 5b). Importantly, and in contrast to cytoBqsS alone, the binding of Fe^2+^ to intact *Pa*BqsS substantially increased its basal ATPase activity with strong significance (Fig. 5b; 15.35 ± 1.10 μmol ATP×mg protein^-1^×min^-1^; *P* < 0.001). Dramatically, the presence of other transition metals (including Fe^3+^) as well as Ca^2+^ all failed to elicit this effect (Fig. 5b). These observations show that although *Pa*BqsS has some inherent ATPase activity in the absence of an external stimulus, the presence of Fe^2+^ (and only Fe^2+^) dramatically and significantly increases this activity, corroborating previous transcriptomic data.^30^ These results are wholly consistent with our hypothesis that the binding of Fe^2+^ within the extracytoplasmic domain is communicated across the lipid bilayer to affect HK function and downstream signaling, and such a result is shown here with an intact, metal-sensing HK for the first time.

### Site-directed mutagenesis reveals key residues for Fe^2+^ sensing and signal transduction

Finally, to determine whether modifications in the periplasmic domain affected the ATPase activity of *Pa*BqsS, we tested our purified variants for activity in the presence and absence of Fe^2+^ and compared these results to the WT protein (Fig. 5d). To establish basal activity levels, we first measured the ATPase activity of the E45A, E48A, and N49A variants of *Pa*BqsS in the absence of Fe^2+^ and found them to be comparable to the WT protein (Fig. 5d): E45A exhibited a basal specific activity of 6.22 ± 0.89 μmol ATP×mg protein^-1^×min^-1^, E48A exhibited a basal specific activity of 6.26 ± 0.60 μmol ATP×mg protein^-1^×min^-1^, and N49A exhibited a basal specific activity of 6.03 ± 0.89 μmol ATP×mg protein^-1^×min^-1^. These results confirm that modification of these periplasmic residues do not directly affect the ability of the cytoplasmic DHp and CA domains of *Pa*BqsS to bind and to hydrolyze ATP at basal levels. We then tested the ability of these variant proteins to be stimulated by Fe^2+^. For the E45A variant in which the Fe^2+^ binding stoichiometry of this variant was unaffected but its binding geometry was modified (*vide supra*), we observe no statistically-significant difference between the basal ATPase activity of this variant and its Fe^2+^-stimulated ATPase activity (5.63 ± 0.47 μmol ATP×mg protein^-1^×min^-1^; Fig. 5d). These data indicate that although Fe^2+^ binds to this variant (albeit in a different geometry), the modified coordination of Fe^2+^ in this variant fails to stimulate *Pa*BqsS activity, demonstrating that E45 plays a role in maintaining proper metal coordination. Unsurprisingly, for the E48A variant that no longer binds Fe^2+^ (*vide supra*), we observed no statistically-significant difference between the basal ATPase activity of this variant and its Fe^2+^-stimulated ATPase activity (6.17 ± 0.76 μmol ATP×mg protein^-1×^min^-1^; Fig. 5d). Interestingly, for the N49A variant in which the Fe^2+^ binding stoichiometry and the Fe^2+^ coordination sphere of this variant were both unaffected (*vide supra*), we observed no statistically-significant difference between the basal ATPase activity of this variant and its Fe^2+^-stimulated ATPase activity (5.92 ± 0.30 μmol ATP×mg protein^-1^×min^-1^; Fig. 5d). As Asn^49^ is not directly involved in coordinating the Fe^2+^ ion, we believe this result points to an uncoupling of Fe^2+^ sensing and ATPase activity stimulation, implicating this residue in the signal transduction pathway (likely through H-bonding), which we did not initially expect. Thus, these data reveal several critical residues within the periplasmic domain of *Pa*BqsS that are necessary for Fe^2+^ coordination and signal transduction in this membrane HK.

## DISCUSSION

The sensing of transition metals is important for the survival and the virulence of infectious prokaryotes, as these micronutrients are needed for normal bacterial function and are also leveraged to establish infection, especially within the mammalian host.^11,12,13^ A tight control of transition metal concentration is critical to pathogens, as too little metal disrupts general metabolic function, while too much metal can be toxic to the cell.^9,59,60^ In order to control this balance, bacteria employ TCSs that are able to respond to extracytosolic concentrations of numerous cations, including the most important transition metal ions (Mn^2+^, Fe^2+/3+^, Cu^+^/Ag^+^, Zn^2+^). Once sensed, the commonly-accepted paradigm posits that a signal is passed through the HK, across the membrane, and ultimately to a response regulator (RR) that is phosphorylated and then targets genes involved in homeostasis and virulence (among others) for transcriptional regulation (Fig. 6).^9^ In pathogenic prokaryotes, the ability to sense and to respond to extracytoplasmic iron is particularly important due to its intimate involvement in critical bacterial metabolic processes^46,61–63^, its low abundance in a mammalian host due to nutritional immunity,^64,65^ and its high toxicity if misregulated.^64,66,67^ Moreover, for many pathogens such as *P. aeruginosa*, the success of infection and survival both depend on multiple forms of iron, as this transition metal also promotes formation of biofilms and exacerbates antibiotic resistance.^68–72^ Ferrous iron (Fe^2+^) is of particular interest to a facultative anaerobic pathogen such as *P. aeruginosa* when colonizing within hypoxic and/or acidic niches present in the mammalian host, such as parts of the stomach, the intestines, within lungs of patients suffering from cystic fibrosis, and inside microaerobic biofilms.^37,65,68,69^ However, how a bacterium such as *P. aeruginosa* senses Fe^2+^ in these environments has remained unclear.

**Figure 6.**
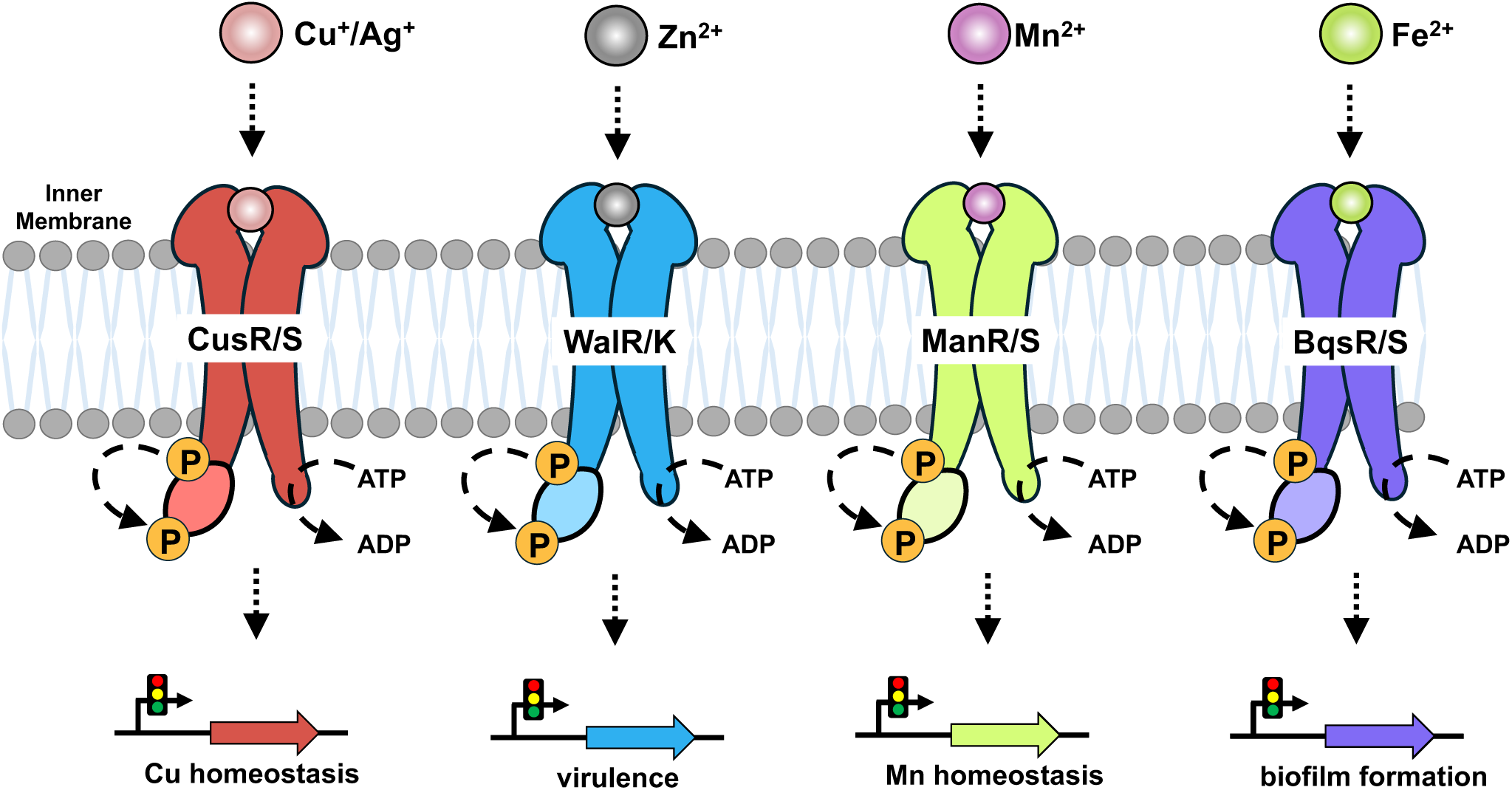
BqsRS complements the functions of other bacterial metal-sensing two-component signal transduction systems (TCSs). Bacterial metal-sensing TCSs that have been partially characterize include: CusRS (red) that senses Cu^+^/Ag^+^ and regulates the expression of Cu efflux transporters and redox enzymes that are responsible for alleviating Cu toxicity through its response regulator (RR); WalRK (blue) that senses Zn^2+^ and regulates the expression of various genes important for virulence, biofilm formation, and cell viability through its RR; and ManRS (green) that senses Mn^2+^ and regulates the expression of genes involved in Mn homeostatsis through its RR. Previous work has shown that BqsRS (purple) regulates the expression of genes involved in biofilm formation and antibiotic resistance, and this work demonstrates that BqsS directly binds and senses Fe^2+^ selectively and responds to the presence of Fe^2+^ by upregulating ATPase activity, representing the first time an intact membrane HK has been show to make a stable and functional interaction with its cognate metal *in vitro*.

To gain a better understanding of how a pathogenic bacterium uses a TCS to sense extracytoplasmic Fe^2+^, we expressed, solubilized, purified, and characterized the intact membrane sensor HK of the BqsRS system, an important biofilm-regulating TCS originally identified in *P. aeruginosa*. We first established that *Pa*BqsS binds a single Fe^2+^ ion at a dimeric interface, and using mutagenic and biophysical approaches, we characterized the composition of this metal-binding site. While we initially hypothesized that an ExxE motif would be involved in coordination of Fe^2+^ due to its conservation among other iron sensing, transport, and storage proteins,^46–49^ we surprisingly found that only the latter Glu residue within the ExxE motif (Glu^48^) was essential for Fe^2+^ binding, while the former Glu residue (Glu^45^) appears to play an auxiliary role in maintaining the correct Fe^2+^ geometry in the periplasmic metal-binding domain. These results represent the first time that an intact, transition-metal sensing membrane HK has been characterized in the presence of its cognate metal, and these findings have interesting implications for the coordination chemistry utilized by other putative Fe-sensing two-component systems that may bind Fe^2+^, Fe^3+^, or both.^50–54^ Moreover, we demonstrated that the binding of Fe^2+^ to *Pa*BqsS has functional ramifications and that disruption within the membrane HK periplasmic domain can uncouple Fe^2+^-based ATPase stimulation. Specifically, when tested against the absence of metal, intact *Pa*BqsS showed a dramatic and significant increase in ATP hydrolysis in the presence of Fe^2+^ that was not observed when just the cytosolic DHp and CA domains of *Pa*BqsS were tested. When investigated against a panel of other metal cations (including both Fe^3+^ and Ca^2+^), this increase in ATP hydrolysis was only observed in the presence of Fe^2+^, demonstrating specificity of both metal ion identity as well as metal ion oxidation state. Importantly, loss of either key residues that coordinate the Fe^2+^ ion (*e.g.*, Glu^45^ and/or Glu^48^) or residues involved in the signal transduction pathway (*e.g.*, Asn^49^) uncouple Fe^2+^ sensing from HK ATPase activation, revealing a direct link between the periplasmic metal-sensing domain and the cytosolic ATPase domain. These consequential observations demonstrate for the first time that the binding of metal within the periplasmic domain of a metal-sensing membrane HK directly functions as a signal that is transmitted to the cytosolic domain of the HK that hydrolyzes ATP and ultimately transmits the phosphate moiety to the RR, significantly advancing our understanding of the structure-function paradigm of a bacterial metal-sensing membrane HK.

These results have important implications for the function of BqsRS in *P. aeruginosa* as well as in other pathogenic bacteria, which could be leveraged for future developments. Previous results have indicated that the BqsRS system may sense Ca^2+^ in addition to Fe^2+^, and that Ca^2+^ sensing via BqsRS may lead to differential downstream gene regulation (thus resulting in BqsRS also being known as CarRS).^31^ Our results here show that Ca^2+^ has no effect on the ATPase activity of *Pa*BqsS directly and that *Pa*BqsS is sensitive to Fe^2+^ alone. Thus, the previously observed Ca^2+^-related effects on the BqsRS/CarRS system^31^ are unlikely to be related to the ability of BqsS to bind and to sense Ca^2+^ itself; however, a protein within the larger *bqsPQRS* operon (locus PA2656-PA2659 in PAO1) or elsewhere within *P. aeruginosa* may instead function to sense Ca^2+^ in this manner.^30^ The intertwined relationship between *bqs* expression and the presence of both Fe^2+^ and Ca^2+^ is perhaps unsurprising, as both of these metals are present at elevated concentrations in the cystic fibrosis lung environment, a common habitat for *P. aeruginosa*, and *bqsS* transcripts have been detected in cystic fibrosis sputum samples.^32–38^ As the BqsRS system regulates rhamnolipid production (an important component of biofilms in *P. aeruginosa*),^73^ the presence of Fe^2+^ could target BqsS directly to regulate biofilm formation via BqsR, while Ca^2+^ may play a role in biofilm dispersal by controlling the *bqs* operon via a separate regulator. As other bacteria (e.g., *Vibrio cholerae, Enterobacter clocae*) also contain BqsRS/CarRS homologs, understanding the intricate but enigmatic crosstalk between Fe^2+^, Ca^2+^, biofilm formation, and biofilm dispersal mediated by this TCS warrants further investigation. Moreover, our results suggest that this system could be a rational target for therapeutics. For example, synthetic inorganic compounds designed to bind to Glu^48^ specifically within the periplasmic domain could be used to stimulate BqsS/CarS and to engage downstream biofilm dispersal, prematurely exposing pathogens to antibiotics in a controlled fashion. In contrast, synthetic small molecules designed to target and to disrupt Asn^49^ specifically within the periplasmic domain could uncouple stimulation of BqsS/CarS, limiting biofilm dispersal and preventing downstream infection. Regardless of the avenue sought, such an approach would require high-resolution structural information on either intact or truncated BqsS/CarS, representing a major future focus on this critical TCS.

## Abbreviations

CA: catalytic ATP-binding domain;
CMC: critical micellular concentration;
CV: column volume;
DHp: dimerization and His phosphotransfer domain;
EXAFS: extended X-ray absorption fine structure;
FI: fit index;
GDN: glyco-diosgenin;
IMAC: immobilized metal affinity chromatography;
IPTG: isopropyl β-D-1-thiogalactopyranoside;
LB: Luria broth;
HK: membrane sensor His kinase;
MWCO: molecular weight cutoff;
OD: optical density;
PMSF: phenylmethylsulfonyl fluoride;
RNS: reactive nitrogen species;
ROS: reactive oxygen species;
RR: response regulator;
SEC: size-exclusion chromatography;
TB: terrific broth;
TEV: tobacco etch virus;
TCS: two-component signal transduction system;
XANES: X-ray absorption near-edge structure;
XAS: X-ray absorption spectroscopy.

## Acknowledgements

This work was supported by NIH-NIGMS grant R35 GM133497 (A.T.S.), NIH-NIGMS grant T32 GM158458 (A.T.S. and A.P.), HHMI Gilliam Fellowship GT15765 (A.P.), and the Murdock Trust Swanson Promise for Scientific Research Award (K.N.C.). Use of the Stanford Synchrotron Radiation Lightsource, SLAC National Accelerator Laboratory, is supported by the U.S. Department of Energy, Office of Science, Office of Basic Energy Sciences under Contract No. DE-AC02-76SF00515. The SSRL Structural Molecular Biology Program is supported by the DOE Office of Biological and Environmental Research, and by the National Institutes of Health, National Institute of General Medical Sciences (P30GM133894). The contents of this publication are solely the responsibility of the authors and do not necessarily represent the official views of NIGMS or NIH. Sequence searches utilized both database and analysis functions of the Universal Protein Resource (UniProt) Knowledgebase and Reference Clusters (http://www.uniprot.org) and the National Center for Biotechnology Information (http://www.ncbi.nlm.nih.gov/).

## Author contributions

A.P., K.N.C., and A.T.S. designed the research; A.P., C.I., and K.N.C. performed the research; A.P., K.N.C., and A.T.S. analyzed the data; and A.P., K.N.C., and A.T.S. wrote and edited the paper.

## Additional information

### Supplementary information

Supplementary information accompanies this paper that includes supplementary figures S1-S8, supplementary tables 1-2, as well as any additional experimental details, materials, and methods.

## METHODS

### Materials

All codon-optimized genes used in this study were synthesized and verified by GenScript. Materials used for buffer preparation, protein expression, and protein purification were purchased from standard commercial suppliers and were used as received. Detergents and Na_2A_TP were purchased from Anatrace, MilliporeSigma, and/or RPI, stored at -20 °C, and used as received. Primers for all site-directed mutagenesis experiments were synthesized and verified by Integrated DNA Technologies.

### Protein expression and purification

Codon-optimized DNA blocks encoding for *Pseudomonas aeruginosa* strain PAO1 BqsS (Uniprot identifier Q9I0I2) either intact (amino acids 1-445, *Pa*BqsS), the RAxxA periplasmic variant (R^44^A^45^xxA^48^), or the cytosolic domain containing both the catalytic ATP-binding and dimerization-His phosphotransfer domains (amino acids 210-440, cyto-*Pa*BqsS), all modified to encode for an additional C-terminal Tobacco Etch Virus (TEV) protease site (ENLYFQS) were commercially synthesized by Genscript. These DNA sequences were then subcloned into the pET-21a(+) expression vector such that the resulting expression construct was read in-frame with a terminating C-terminal (His)_6 t_ag. The expression plasmid was electroporated into electrocompetent *Escherichia coli* BL21 (DE3) cells, spread onto Luria Broth (LB) agar plates supplemented with 0.1 mg/mL ampicillin (final), and grown overnight at 37 °C.

Large-scale expression of intact *Pa*BqsS and variant constructs (E44A, E48A, and E44A/E48A; *vide infra*) was accomplished in 6 baffled flasks each containing 1 L sterile terrific broth (TB) media supplemented with 0.1 mg/mL ampicillin (final) and inoculated with a pre-culture derived from a single bacterial colony. Cells were grown at 37 °C with shaking of 200 RPM until the optical density (OD) at 600 nm (OD_600)_ reached *ca.* 1.2-1.5, at which point cell cultures were cold-shocked for 2 hr at 4 °C. After cold shocking, another bolus of ampicillin was added to each flask, and protein production was then induced with 1 mM (final) isopropyl β-D-1-thiogalactopyranoside (IPTG). Cells were shaken overnight at 200 RPM and 18 °C. After 16-20 hr incubation, cells were harvested by centrifugation at 5000 ×*g* for 12 min at 4 °C, and cell pellets were resuspended in a cellular resuspension buffer (25 mM Tris, pH 8.0, 100 mM sucrose). Immediately prior to cell lysis, phenylmethylsulfonyl fluoride (PMSF, *ca.* 50-100 mg) was added. Cells were then kept cold in an ice bath while being disrupted via sonication (80 % maximal amplitude, 30 s on pulse, 30 s off pulse, 12 min total time on). Cellular debris was cleared by centrifugation at 10,000 ×*g* and 4 °C for 1 hr. The supernatant was decanted and used to prepare crude cellular membranes by ultracentrifugation at 160,000 ×*g* and 4°C for 1 hr. These crude membranes were washed with resuspension buffer, homogenized with a Dounce homogenizer, and ultracentrifuged again at 160,000 ×*g* and 4°C for 1 hr to pellet purified membranes. Purified membranes were then diluted with resuspension buffer and homogenized one final time with a Dounce homogenizer. Solubilization of intact *Pa*BqsS and variants thereof was accomplished by dropwise addition of a stock solution of 10% (w/v) detergent in 4M NaCl until a final concentration of 1% (w/v) detergent and of 500 mM NaCl was reached. During detergent and salt addition, vigorous stirring occurred at room temperature (RT), and the solubilization solution was allowed to continue for *ca.* 3 h at RT. The solution was then clarified by ultracentrifugation at 160,000 ×*g* and 4°C for 1 hr, and the supernatant was then purified via immobilized metal affinity chromatography (IMAC). Briefly, the supernatant containing the Fos-Chol-14-solubilized *Pa*BqsS was applied to a prepacked 5 mL HisTrap HP column (Cytiva). The column was then washed with 5 column volumes (CVs) of wash buffer (50 mM Tris, pH 8.0, 300 mM NaCl, 1 mM TCEP, 10 % (v/v) glycerol, and 0.05% (w/v) Fos-Chol-14) and protein was then eluted stepwise with 7.0%, 16.7%, 50.0%, and 100.0% (all v/v) respective linear gradients of elution buffer (the same composition of the wash buffer plus 300 mM imidazole). Fractions containing purified *Pa*BqsS were pooled and concentrated using a 100 kDa molecular weight cutoff (MWCO) Amicon spin concentrator. The protein was then combined with *ca.* 20 μg of TEV protease per *ca.* 1 mg *Pa*BqsS and allowed to rock overnight at 4°C in order to cleave the (His)_6 t_ag. Following tag cleavage, another IMAC run was performed and the now cleaved *Pa*BqsS was collected from the flow-through and concentrated using a 100 kDa MWCO Amicon spin concentrator. Cleaved *Pa*BqsS was then applied to a HiLoad 16/600 Superose 6 pg preparative SEC column (Cytiva) pre-equilibrated with SEC buffer (50 mM Tris, pH 7.5, 100 mM NaCl, 1 mM TCEP, 5 % (v/v) glycerol, and 0.02% (w/v) Fos-Chol-14) and eluted isocratically. Fractions containing homogenous protein were collected, quantified, and analyzed using 15% SDS-PAGE and western blotting.

Expression and purification of cyto*Pa*BqsS was accomplished differently than the intact membrane protein. Large-scale expression of cyto*Pa*BqsS was accomplished in 6 baffled flasks each containing 1 L sterile LB supplemented with 0.1 mg/mL ampicillin (final) and inoculated with a pre-culture derived from a single bacterial colony. Cells were shaken at 200 RPM until the OD_600 r_eached *ca*. 0.6-0.8. Cell cultures were then cold-shocked for 1.5-2 hours at 4 °C before being induced with 1 mM (final) isopropyl β-D-1-thiogalactopyranoside (IPTG) and incubated at 18 °C overnight with continued shaking. Cells were then harvested by centrifugation for 12 min at 5000×*g* and 4 °C and resuspended with a cellular resuspension buffer (50 mM Tris, pH 7.5, 300 mM NaCl, 10 % (v/v) glycerol, 1 mM TCEP). Immediately prior to cell lysis, phenylmethylsulfonyl fluoride (PMSF, *ca.* 50-100 mg) was added. Cells were then kept cold in an ice bath while being disrupted via sonication (80 % maximal amplitude, 30 s on pulse, 30 s off pulse, 12 min total time on). The resulting lysate was clarified by ultracentrifugation at 160,000×*g*, 4 °C for 1 hr. The clarified lysate containing was then purified via immobilized metal affinity chromatography (IMAC) and size-exclusion chromatography (SEC). Briefly, the lysate was injected onto a prepacked 5 mL HisTrap HP column (Cytiva). The column was then washed with 5 column volumes (CVs) of wash buffer (50 mM Tris, pH 7.5, 300 mM NaCl, 10 % (v/v) glycerol, 1 mM TCEP) and eluted step-wise with 7.0 %, 16.7 %, 50.0 %, and 100.0 % (all v/v) respective linear gradients of elution buffer (the same composition of the wash buffer plus 300 mM imidazole). Fractions containing intact cyto*Pa*BqsS were pooled and concentrated using a 10 kDa MWCO Amicon spin concentrator. The protein was then buffer exchanged into a TEV cleavage buffer (50 mM Tris, pH 8.0, 200 mM NaCl, 5 % (v/v) glycerol, 1 mM TCEP, 0.5 mM EDTA) by repeated concentrations and dilutions in the same spin concentrator. A mixture was then made comprising *ca*. 20 μg TEV protease per *ca*. 1 mg protein and allowed to rock at 4 °C overnight. Following tag cleavage, the protein sample was applied to a HiLoad 10/300 GL Superdex 75 preparative SEC column (Cytiva) pre-equilibrated with SEC buffer (50 mM Tris, pH 7.5, 100 mM NaCl, 5 % (v/v) glycerol, 1 mM TCEP) and eluted isocratically. Fractions containing homogenous protein were collected, quantified, and analyzed using 15 % SDS-PAGE and western blotting.

### Mass photometry of *Pa*BqsS

The distribution of particle size in the purified samples of *Pa*BqsS was determined via mass photometry using a Refyn TwoMP instrument. First, a linear calibration curve (R^2^ = 0.999) was created using β-amylase and thyroglobulin (58 kDa, 113 kDa, 228 kDa, and 670 kDa, respectively). Purified *Pa*BqsS was deposited onto a glass slide at a concentration of *ca.* 100 nM, then rapidly diluted to a 1:100 (v:v) dilution in SEC buffer lacking detergent. Recordings were analyzed using the DiscoverMP software, and the molecular weights of *Pa*BqsS and Fos-Choline-14 micelles were interpolated using the linear standard curve.

### Site-directed mutagenesis

Mutations to the intact *Pa*BqsS expression plasmid were accomplished by the QuikChange Lightning Multi Site-Directed Mutagenesis kit (Agilent) in order to encode for the following protein variants with modifications in the periplasmic metal-binding motif: E45A, E48A, N49A, and E45A/E48A. The following primers (Table S2) were used to introduce the mutated bases (underlined): E45A; forward 5’-GTTCTCCGCTTCC<u>G</u>CACGCAGGTTGCC-3’ reverse: 5’-GGCAACCTGCGTG<u>C</u>GGAAGCGGAGAAC-3’; E48A; forward: 5’-

CAACCAGCAGGTTCGCCGCTTCCTCACGC-3’ reverse: 5’-

GCGTGAGGAAGCGGCGAACCTGCTGGTTG-3’; N49A; forward: 5’-

ATCGCAACCAGCAGGGCCTCCGCTTCCTCACG -3’; reverse: 5’-

CGTGAGGAAGCGGAG<u>GC</u>CCTGCTGGTTGCGAT -3’. The presence of each mutation was verified by nanopore DNA sequencing.

### Metal-binding assays

All (His)_6-_cleaved *Pa*BqsS constructs and its variants were tested for their ability to bind Fe^2+^. Briefly, each construct was degassed and brought into an anoxic chamber containing an N_2/_H_2 a_tmosphere and operating at < 5 ppm O_2._ After overnight equilibration with the N_2/_H_2 a_tmosphere, each protein solution was incubated at 6 °C for 15 min with excess (10 mol eq.) of Fe^2+^. After the incubation period, the protein-metal solution was buffer exchanged into SEC buffer (50 mM Tris, pH 7.5, 100 mM NaCl, 1 mM TCEP, 5 % (v/v) glycerol, and 0.02% (w/v) Fos-Chol-14) through extensive rounds of concentration and dilution using a 0.5 mL 100 kDa MWCO spin concentrator. To quantitate the amount of Fe^2+^ bound, a modified version of the ferrozine assay^74,75^ as previously described.^76^ Briefly, the protein sample was precipitated with 5 M trichloroacetic acid, the solution centrifuged, and the supernatant was collected after being neutralized by saturated ammonium acetate. Freshly prepared excess ascorbic acid and 0.30 mM ferrozine (final concentration) were added to the supernatant. The absorbance at 562 nm was used to quantitate iron content assuming a Fe^2+^-ferrozine complex with an extinction coefficient (ε_562)_ of ≈ 28 mmol L^-1^ cm^-1^.^74,75^ All iron content analysis was done in triplicate.

### X-ray absorption spectroscopy (XAS)

Samples containing *Pa*BqsS and its variants were metal-loaded anoxically, were concentrated to *ca.* 1-2 mM Fe^2+^ (final concentration), were mixed anoxically with ethylene glycol (20% (v/v) final), and were aliquoted anoxically into Lucite cells wrapped with Mylar tape, flash-frozen in liquid N_2 a_nd stored at -80 °C until data collection. X-ray absorption spectroscopy (XAS) was performed on beamlines 7-3 and 9-3 at the Stanford Synchrotron Radiation Lightsource (Menlo Park, CA), and samples in replicates were collected when possible. Extended X-ray absorption fine structure (EXAFS) of Fe (7210 eV) was measured using a Si 220 monochromator with crystal orientation φ = 90°. Samples were measured as frozen aqueous glasses at 15 K, and the X-ray absorbance was detected as Kα fluorescence using either a 100-element (beamline 9-3) or 30-element (beamline 7-3) Ge array detector. A Z-1 metal oxide filter (Mn) and slit assembly were placed in front of the detector to attenuate the elastic scatter peak. A sample-appropriate number of scans of a buffer blank were measured at the absorption edge and subtracted from the raw data to produce a flat pre-edge and eliminate residual Mn Kβ fluorescence of the metal oxide filter. Energy calibration was achieved by placing a Fe metal foil between the second and third ionization chamber. Data reduction and background subtraction were performed using EXAFSPAK (Microsoft Windows version).^77^ The data from each detector channel were inspected for drop outs and glitches before being included into the final average. EXAFS simulation was carried out using the program EXCURVE (version 9.2) as previously described.^77–79^ The quality of the fits was determined using the least-squares fitting parameter, *F*, which referred to as the fit index (FI) and is defined as:

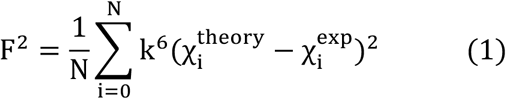

### Structural modeling

All AlphaFold models were acquired from the predicted AlphaFold Protein Structure Database^80^ or modeled *de novo* using the AlphaFold3 structure prediction server.^81^ In order to obtain the predicted structures as well as any aligned error, the following AlphaFold model IDs or UniProt IDs were used, respectively: AF-Q9I0I2-F1-v4 and Q9I0I2.

### ATP hydrolysis assays

The rate of ATP hydrolysis as a proxy for kinase activity was determined using a modified form of the malachite green assay that has been adapted for membrane proteins in the presence of detergents.^58^ Briefly, stocks of WT *Pa*BqsS were diluted to 0.80-3.0 mg/mL (final) using SEC buffer. Either anoxic stock solutions of (NH_4)2F_e(SO_4)2 (_added to anoxic WT *Pa*BqsS or cytoBqsS) or oxic stock solutions of ZnCl_2,_ CoCl_2,_ CaCl_2,_ MgCl_2,_ CuCl_2,_ MnCl_2,_ and FeCl_3 r_anging from 0.5-100 mM (final concentration) were added to *Pa*BqsS or cytoBqsS to stimulate activity. Stock solutions of substrate (Na_2A_TP, 0-1.60 mM final concentration) and Mg^2+^ (5.0 mM final concentration) were added to initiate the reaction. Anoxic assays were carried out within an anoxic chamber (*vide supra*) and all reactions were run at 37 °C with 300 RPM shaking for 0-30 min, depending on the construct. The reaction was then immediately halted by addition of a 3:1 (v:v) ratio of working solution (1.05 % (w/v) ammonium molybdate tetrahydrate, 0.0338 % (w/v) malachite green carbinol, 1.0 M HCl) to the protein assay solution. After addition of 100 μL 34 % (w/v) sodium citrate, the absorbance at 660 nm of the enzyme-containing solution was measured using Cary 60 UV-Vis spectrophotometer (Agilent), and [P_i]_ was interpolated using a standard curve. Specific activity data are corrected against background hydrolysis of Na_2A_TP in the absence of enzyme ± metal.

## For Table of Contents Use Only

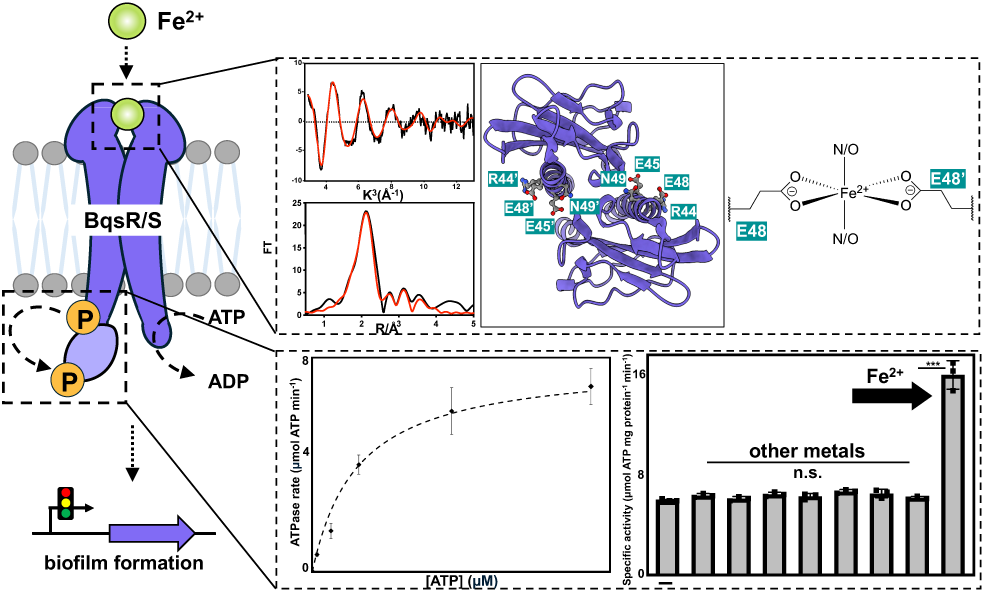

