## Supporting Information for "The *Pseudomonas aeruginosa* Membrane Histidine Kinase BqsS/CarS Directly Senses Environmental Ferrous Iron (Fe^2+^)"

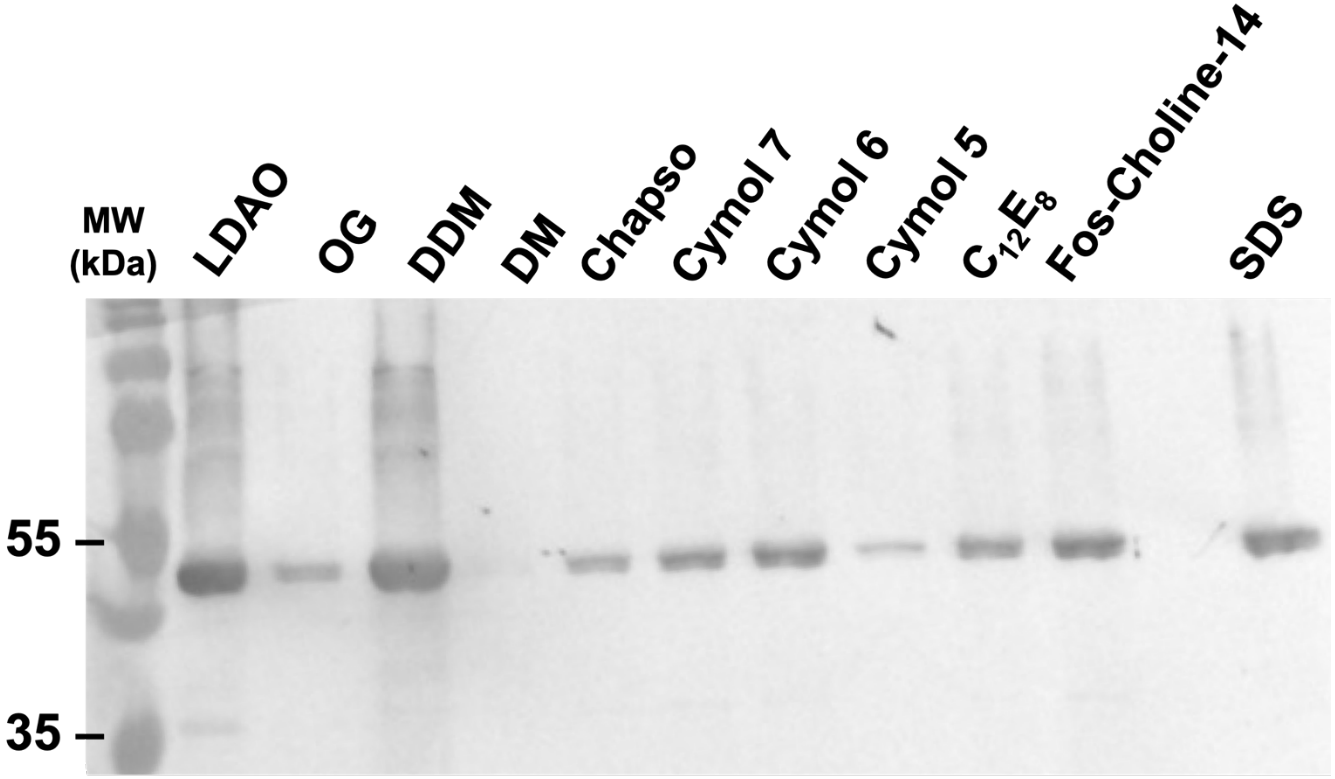


**Figure S1**. Detergent solubilization tests of *P. aeruginosa* BqsS. Normalized western blot analysis of the efficacy of various detergents to solubilize intact *Pa*BqsS (molecular weight of *ca*. 52 kDa). Of the detergents tested, LDAO, DDM, and Fos-Choline-14 showed the highest solubilization efficacy compared to the SDS control (far right lane). Ultimately, Fos-Choline-14 was chosen due to its ability to solubilize *Pa*BqsS stably and homogenously.

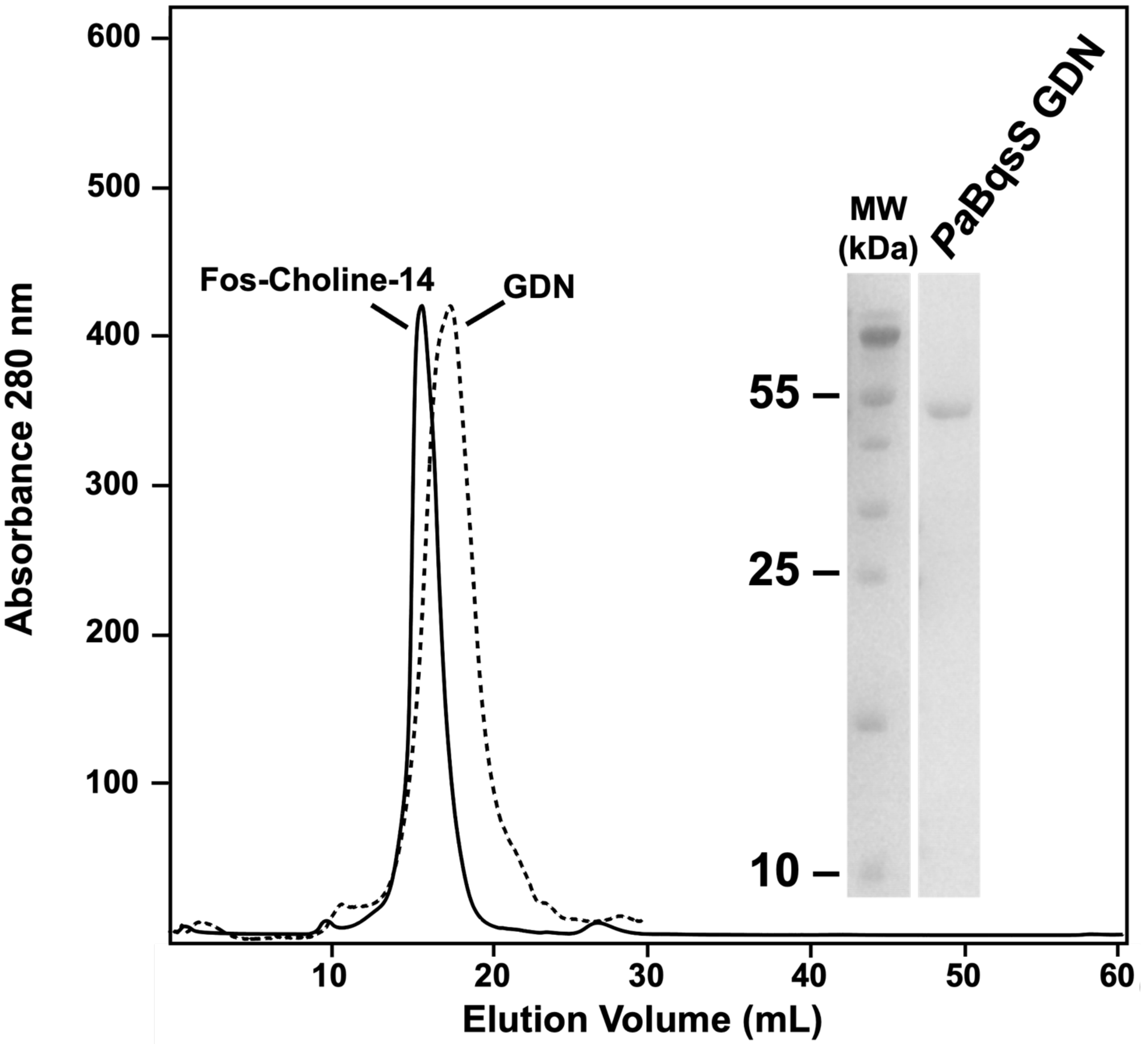


**Figure S2**. Size-exclusion chromatography is most consistent with purified *Pa*BqsS solubilized in Fos-Choline-14 existing as a dimer of dimers (solid), while *Pa*BqsS solubilized in glyco-diosgenin (GDN) is most consistent with a dimeric quaternary structure (dashed). *Inset.* 15 % SDS-PAGE analysis of *Pa*BqsS after solubilization and purification in GDN.

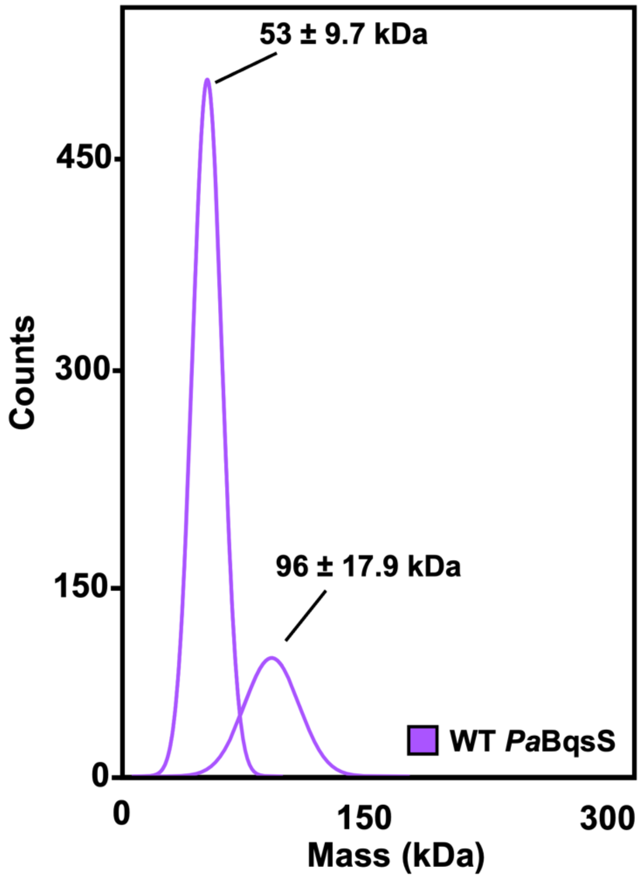


**Figure S3**. Mass profile of WT *Pa*BqsS in Fos-Choline-14 determined by mass photometry. Empty Fos-Choline-14 micelles are observed at *ca*. 53 kDa, while dimeric WT *Pa*BqsS is observed at *ca*. 96 kDa. The molecular weights of the gaussian fits were determined by use of a calibration curve of two standards: β-amylase and thyroglobulin.

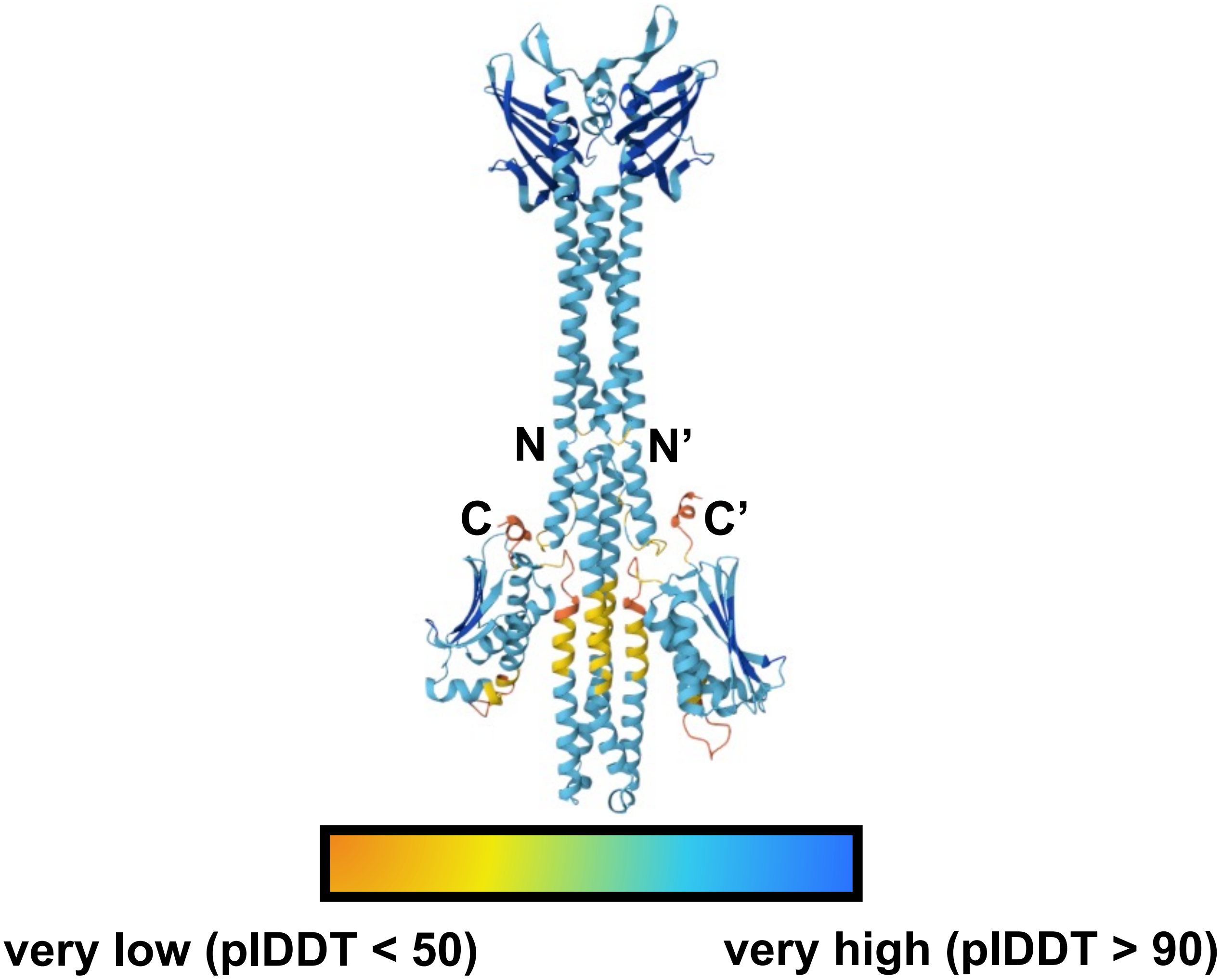


**Figure S4**. AlphaFold model of intact, dimeric *Pa*BqsS color-coded with the per-residue confidence score (plDDT) from very low confidence (orange) to very high confidence (blue).

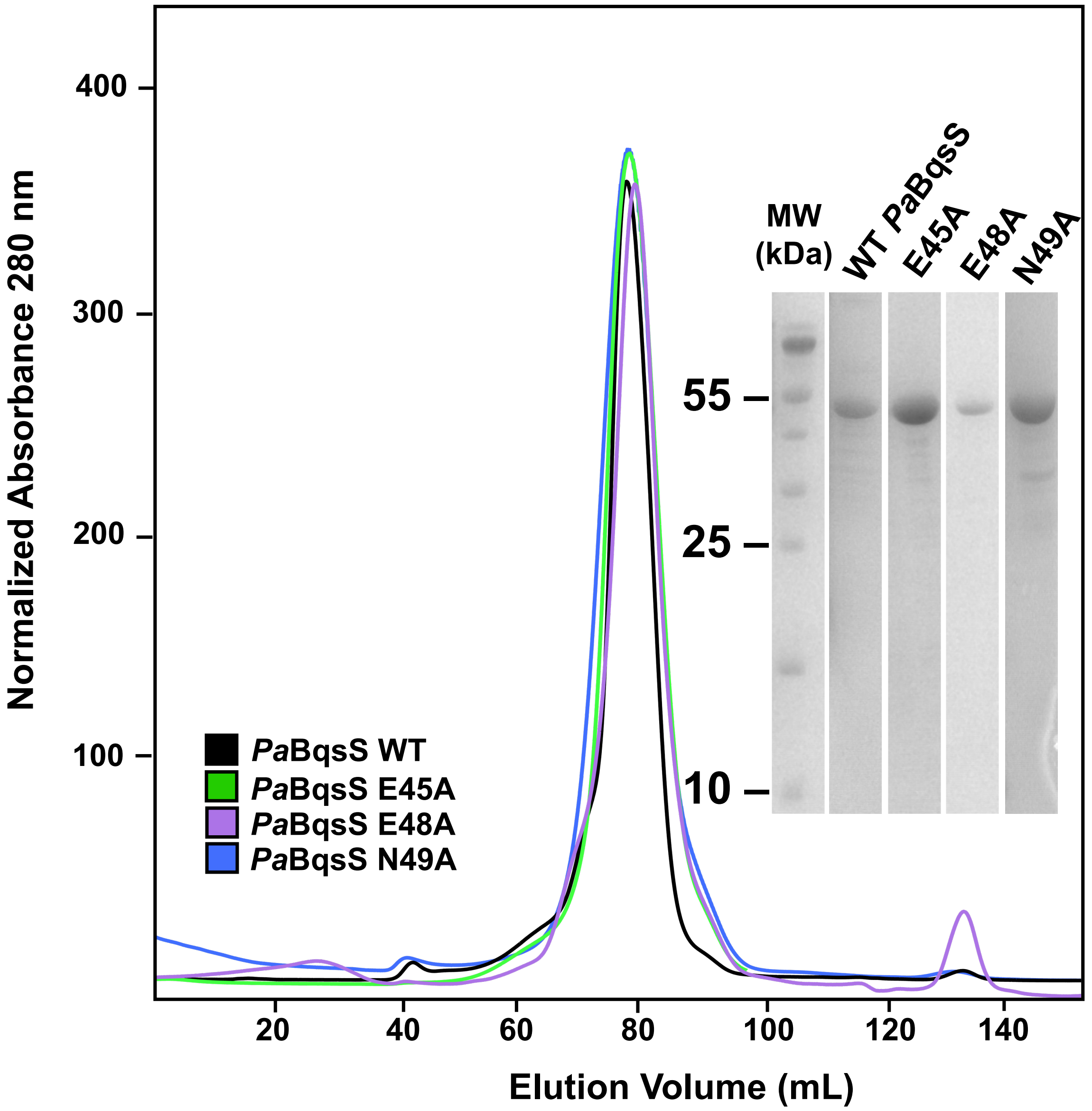


**Figure S5**. WT *Pa*BqsS and *Pa*BqsS variants E45A, E48A, and N49A have similar purities in Fos-Choline-14 (*inset*; 15% SDS-PAGE), and all migrate identically based on size-exclusion chromatography.

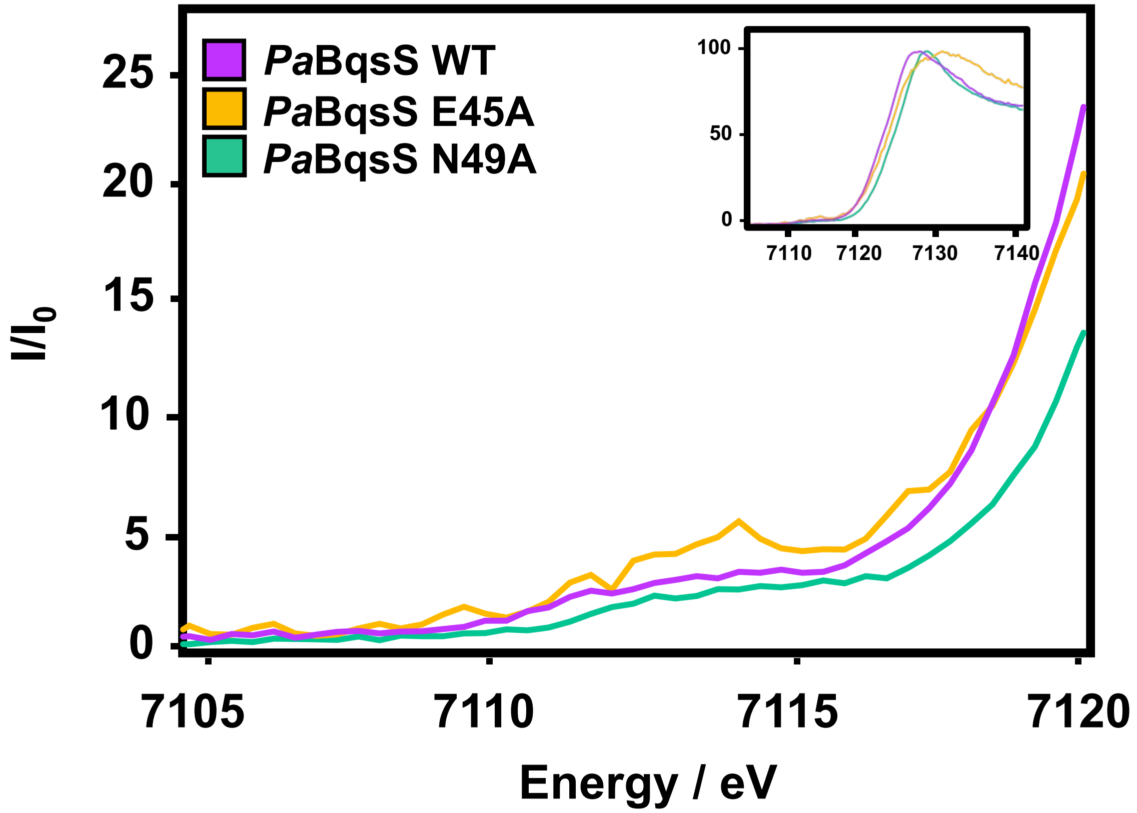


**Figure S6**. The pre-edge features found in the normalized X-ray absorption near-edge structure (XANES) spectrum of intact, $Fe^{2+}$-bound WT *Pa*BqsS (purple), E45A *Pa*BqsS (orange), and N49A *Pa*BqsS (green). *Inset*: full XANES spectra of intact, $Fe^{2+}$-bound WT *Pa*BqsS (purple), E45A *Pa*BqsS (orange), and N49A *Pa*BqsS (green).

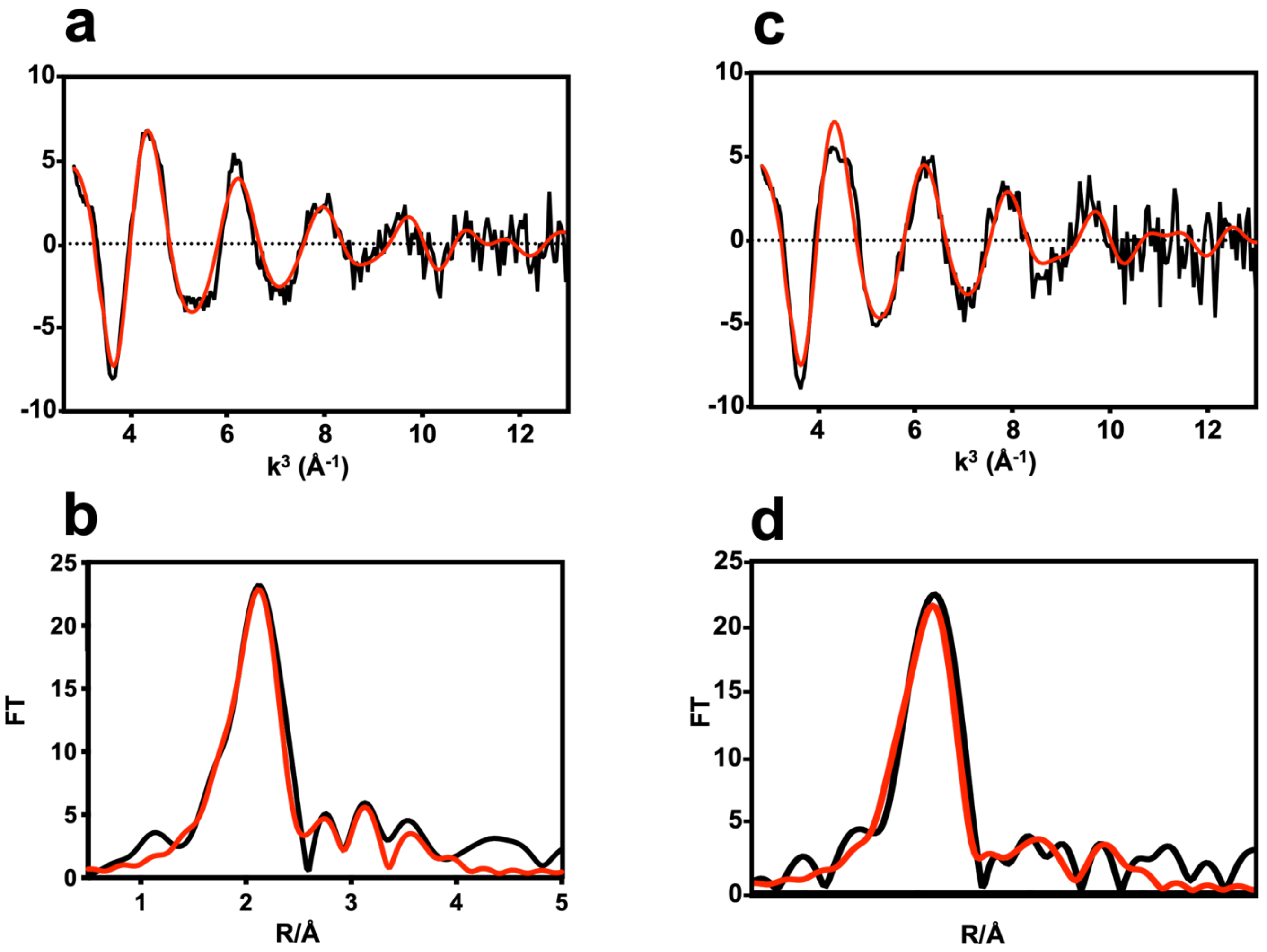


**Figure S7**. The processed WT *Pa*BqsS EXAFS (**a**) and resulting Fourier transformed data (**b**) are consistent with the presence of $Fe^{2+}$ in an octahedral geometry composed of an N/O-rich environment. The processed N49A *Pa*BqsS EXAFS (**c**) and resulting Fourier transformed data (**d**) suggest a nearly identical ligation sphere. The black traces represent the experimental data while the red traces represent the EXCURVE-fitted data.

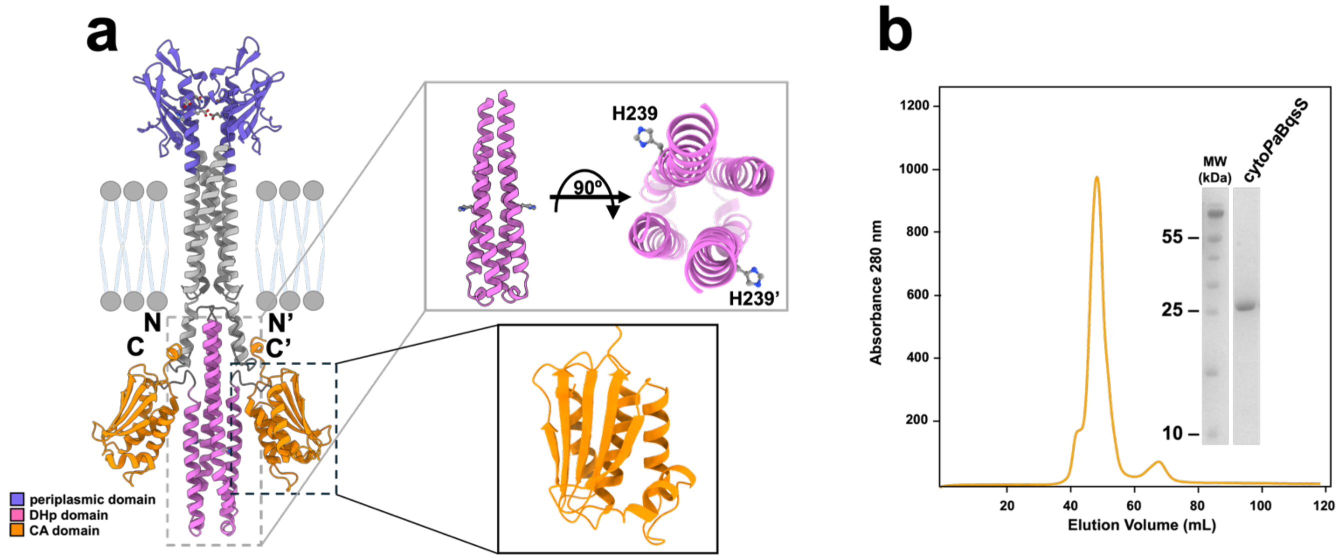


**Figure S8**. Modeling and purification of the cytosolic DHp and CA domains of *Pa*BqsS (cytoBqsS). **a**. The AlphaFold model of intact *Pa*BqsS with the various domains (periplasmic, DHp, CA) color-coded. *Inset*. The cytoplasmic domain of *Pa*BqsS (cytoBqsS) consists of a dimerization and His phosphotransfer (DHp) domain (pink) that houses a conserved phosphate-accepting His residue ($His^{239}$) and a catalytic ATP-binding domain (CA) domain (orange) that catalyzes the transfer of the γ-phosphate from ATP to $His^{239}$. **b**. Size-exclusion chromatogram (SEC) and 15 % SDS-PAGE analysis (*inset*) of cytoBqsS (molecular weight of *ca*. 27 kDa).
